# PI3K/AKT inhibition in tumor propagating cells of DLBCL reverses R-CHOP resistance by destabilizing SOX2

**DOI:** 10.1101/657445

**Authors:** Jianfeng Chen, Xiaowen Ge, Wei Zhang, Peipei Ding, Yiqun Du, Qi Wang, Ling Li, Lan Fang, Yujing Sun, Pingzhao Zhang, Yuzhen Zhou, Long Zhang, Xinyue Lv, Luying Li, Xin Zhang, Qunling Zhang, Kai Xue, Hongyu Gu, Qunying Lei, Jiemin Wong, Weiguo Hu

## Abstract

Drug resistance is a major obstacle for the success of conventional anticancer therapy, and the development of drug resistance is at least partly attributed to tumor propagating cells (TPCs). Up to one-third of diffuse large B cell lymphoma (DLBCL) patients eventually develop resistance to R-CHOP regimen. We found that the TPC proportion was remarkably increased in resistant germinal center B cell-like (GCB) and activated B cell-like (ABC) DLBCL subtypes. Elevated SOX2 was the determinant for resistance development, and SOX2 was phosphorylated by activated PI3K/AKT1 signaling, thus preventing ubiquitin-mediated SOX2 degradation. Furthermore, multiple factors, including BCR, integrins, chemokines and FGFR1/2 signaling, regulated PI3K/AKT1 activation. CDK6 in the GCB subtype and FGFR1/2 in the ABC subtype were SOX2 targets in the PI3K/AKT1 pathway. Chemical inhibition of PI3K/AKT1 in both subtypes, CDK6 in the GCB subtype, and FGFR1/2 in the ABC subtype significantly enhanced the susceptibility of resistant cells to CHO treatment. More importantly, PI3K and FGFR1/2 inhibitors but not a CDK6 inhibitor effectively suppressed the tumor growth of R-CHO-resistant DLBCL cells, most likely by converting TPCs to chemo-sensitive differentiated cells. Therefore, this pro-differentiation therapy against TPCs warrants further study in clinical trials for the treatment of resistant DLBCL.

## Introduction

The success of conventional anticancer therapy is frequently hindered by drug resistance. Such acquired resistance results at least partly from intratumoral heterogeneity that is accounted for the predominant differentiated cancer cells and the tiny subpopulation of tumor propagating cells (TPCs). Although most differentiated cells are eliminated by conventional therapy, TPCs survive, which may drive drug resistance development (*1, 2*). TPCs are characterized by distinct surface proteins, self-renewal, differentiation, quiescence, and a high association with drug resistance, recurrence and metastasis (*2–5*). Therefore, a combination of conventional therapy and TPC-specific therapy is strongly indicated and being pursued (*6, 7*).

SOX2, a pluripotency-associated transcription factor, is critical for embryonic (*8*) and induced pluripotent stem cells (*9*); however, its expression is highly detrimental in at least 25 different cancers (*10*). In contrast with numerous studies focusing on *SOX2* regulation by non-coding RNAs, there have been limited reports concerning transcriptional regulation and post-translational modifications. PI3K/AKT1 signaling is a master regulator not only in tumorigenesis, tumor progression, and drug resistance (*11*) but also in TPC biology (*12*). Interestingly, PI3K/AKT1 may suppress SOX2 ubiquitination via a methylation (K119)-phosphorylation (T118) switch in SOX2, thus stabilizing SOX2 (*13*).

Non-Hodgkin’s lymphoma (NHL) ranks in the top 10 causes of cancer mortality, and diffuse large B cell lymphoma (DLBCL) is the most common subtype (*14*). DLBCL can be subdivided into three distinct cell-of-origin subtypes: germinal center B cell-like (GCB), activated B cell-like (ABC), and 10-20% primary mediastinal B cell lymphoma (PMBL) subtypes (*15*). Although more than half of DLBCL patients can be cured, mainly by R-CHOP (rituximab/R, cyclophosphamide/C, doxorubicin/H, vincristine/O, and prednisone/P) regimens (*16*), up to one-third of patients will eventually develop relapsed/refractory disease (*17*). Our growing understanding of the molecular basis of resistance has led to the development of a large number of novel interventions, however, they are only being tested in phase I or II trials, and no single agent or regimen provides long-term disease control (*18, 19*). Thus, novel therapeutic approaches for relapsed/refractory DLBCL are urgently needed.

Here we found a remarkably elevated proportion of TPCs in resistant DLBCL cells, whose stemness was regulated by the activated PI3K/AKT1/SOX2 axis. Further, PI3K/AKT inhibitor converted TPCs to differentiated tumor cells by reducing SOX2 level, thus preventing the growth of implanted resistant cells when combined with the R-CHOP regimen.

## Methods

### DLBCL tissue samples, cell lines and reagents

We examined the medical history of all DLBCL patients from 2008 to 2015 at Fudan University Shanghai Cancer Center and found a total of 12 patients who simultaneously had both paraffin-embedded tissue samples from the initial visit and from relapse. DLBCL cases were subgrouped into GCB (6 cases) or ABC (6 cases) molecular subtypes based on the Hans immunohistochemistry algorithm. Additional information is provided in the supplemental material.

### Generation and characterization of RCHO-resistant DLBCL cells

We generated RCHO-resistant LY8 (OCI-Ly8) and NU-DUL-1 cells as previously described and details are described in the supplemental material (*20, 21*).

### Aldefluor Assay

ALDH1 is a selectable marker for multiple kinds of normal and cancer stem cells, including hematopoietic stem cells (*22, 23*). Thus, we evaluated tumor propagating cell numbers in hematopoietic malignancies using an ALDEFLUOR™ kit (StemCell Technologies, Vancouver, BC, CA) to detect ALDH1^+^ cells. Details are described in the supplemental material.

### Sphere Formation Assay

We conducted sphere formation assays as previously described and details are reported in the supplemental material (*24*).

### CytoTox-Glo™ Cytotoxicity Assay

We used a CytoTox-Glo™ cytotoxicity assay kit (Promega, Madison, WI) to evaluate cytotoxicity according to the technical bulletin. Details are described in the supplemental material.

### Immunohistochemistry

Tumor tissues derived from patients or animal models were fixed with 4% formalin, embedded in paraffin and sectioned. Immunohistochemistry (IHC) staining was performed on 4 μm paraffin sections and details are provided in the supplemental material.

### Immunoblotting Assay

We performed immunoblotting assays according to the standard protocol, and the related antibodies are shown in Table S1.

### Quantitative Real-time PCR (qRT-PCR)

We performed qRT-PCR as previously described and details are reported in the supplemental material(*24*). The primers for qRT-PCR are listed in Table S2.

### FACS Analysis

Flow cytometric analysis was performed on a Cytomics FC500 MPL instrument (Beckman Coulter, Brea, CA) and analyzed with FlowJo software (Ashland, OR). We performed cell sorting with a MoFlo XDP instrument (Beckman Coulter, Brea, CA). Details are described in the supplemental material.

### EdU Cell Proliferation Assay

The EdU cell proliferation assays were performed using the Cell-Light EdU Apollo643 In Vitro Flow Cytometry Kit (Guangzhou RiboBio Co., Ltd., Guangzhou, China). Details are described in the supplemental material.

### Plasmid Construction and Lentiviral Transduction

The plasmid construction and lentiviral transduction was performed as previously described and details are depicted in the supplemental material(*24*). Information about shRNA oligonucleotide sequences is shown in Table S2.

### Xenograft Model

All the animal experiments were conducted in strict accordance with experimental protocols approved by the Animal Ethics Committee at Shanghai Medical School, Fudan University. Eight-week-old female SCID mice were purchased from Slac Laboratory Animal Center (Shanghai, China) for injection with RCHO-resistant DLBCL cells. The methods of drug delivery based on the clinical usage for one cycle are indicated in Table S3. Tumor growth was monitored by bioluminescence at 50, 70 and 90 days after implantation using an In Vivo MS FX PRO system (Bruker, Billerica, MA). The surviving mice were euthanized and dissected at 120 days after xenografting, and no intraperitoneal tumors were found. Tumor tissues were immediately collected from the moribund mice after euthanatized by CO_2_. Additional information including serial-transplantation for detecting tumor-initiating capacity of RCHO-resistant cells is provided in the supplemental material.

### RNA Sequencing and Bioinformatic Analysis

Total RNA was extracted from LY8-ORI, LY8-R, LY8-CHO, LY8-RCHO, NU-DUL-1, NU-DUL-1-R, NU-DUL-1-CHO, and NU-DUL-1-RCHO cells with TRIzol reagent (Invitrogen, Grand Island, NY). The total RNA from each group from 3 different passages was pooled separately. RNA sequencing (RNA-seq) and bioinformatics analysis were conducted by Shanghai Novelbio Ltd. (*25*). Details are decribed in the supplemental material.

## Results

### Resistance of stem-like resistant DLBCL cells was regulated by SOX2

Antibody-dependent cellular cytotoxicity (ADCC) and complement-dependent cytotoxicity (CDC) are the major mechanisms underlying R therapeutic effect (*26, 27*). Resistance to ADCC may derive from intrinsic features of immune cells of individual patient, such as *FcGRIIIA* polymorphism and CD47 expression (*28, 29*); while resistance to CDC is mediated by a low expression level of CD20 and/or high expression levels of membrane-bound complement regulatory proteins (mCRPs), including CD46, CD55 and especially CD59 (*27, 30*). CHO (cyclophosphamide/C, doxorubicin/H, vincristine/O) suppress tumor cell duplication by inducing DNA damage or binding to tubulin, and prodrug C must be metabolically activated *in vivo* into 4-hydroxycyclophosphamide (4-HC) to exert effect. Therefore, using R plus normal human serum (NHS, as a complement source), 4-HC plus H plus O, or their combination, we prepared resistant LY8 (GCB subtype) or NU-DUL-1 (ABC subtype) DLBCL cells, named R, CHO, and RCHO, respectively (Fig. 1A). The resistance to CDC (Fig. S1A) and to chemotherapy (Fig. S1B) was validated, and cross-resistance to R-mediated CDC and CHO-mediated chemotherapy was observed to a certain degree. Furthermore, the resistant cells exhibited stem-like spheres with varying sizes compared with the original (ORI) cells, and the number of spheres increased gradually in order of R, CHO, and RCHO (Fig. 1B). Although few studies have experimentally demonstrated the presence of TPCs in NHL, they are still proposed to exist (*31, 32*). Thus, we employed ALDH1 (*33*) and side population (*34*) in TPCs of various cancer types and CD34 and CD133 in TPCs of leukemia (*35, 36*) to ascertain the stemness of resistant DLBCL cells. The results showed a significantly elevated proportion of ALDH1^+^, side population, CD34^+^ and CD133^+^cells in the resistant cells (Fig. 1C-F, and Fig. S1C-F), in which the increase of CD34 and CD133 levels were further verified by immunoblotting (Fig. 1G). These TPC markers may group different subpopulation with partially overlapping, for example, a small fraction of cells showed CD133^+^ among ALDH1^+^ cells (Fig. S2) as described previously (*37*). In addition, we also observed that the expression of PAX5, a B cell commitment factor (*38*), reduced in parallel with the resistance degree (Fig. 1G). More importantly, using serial-transplantation assay we revealed that both RCHO-resistant LY8 and NU-DUL-1 cells displayed higher tumor-initiating capacity than original cells (Table 1). Together, these results strongly demonstrated that TPCs were highly enriched in the resistant DLBCL cells.

**Fig. 1.**
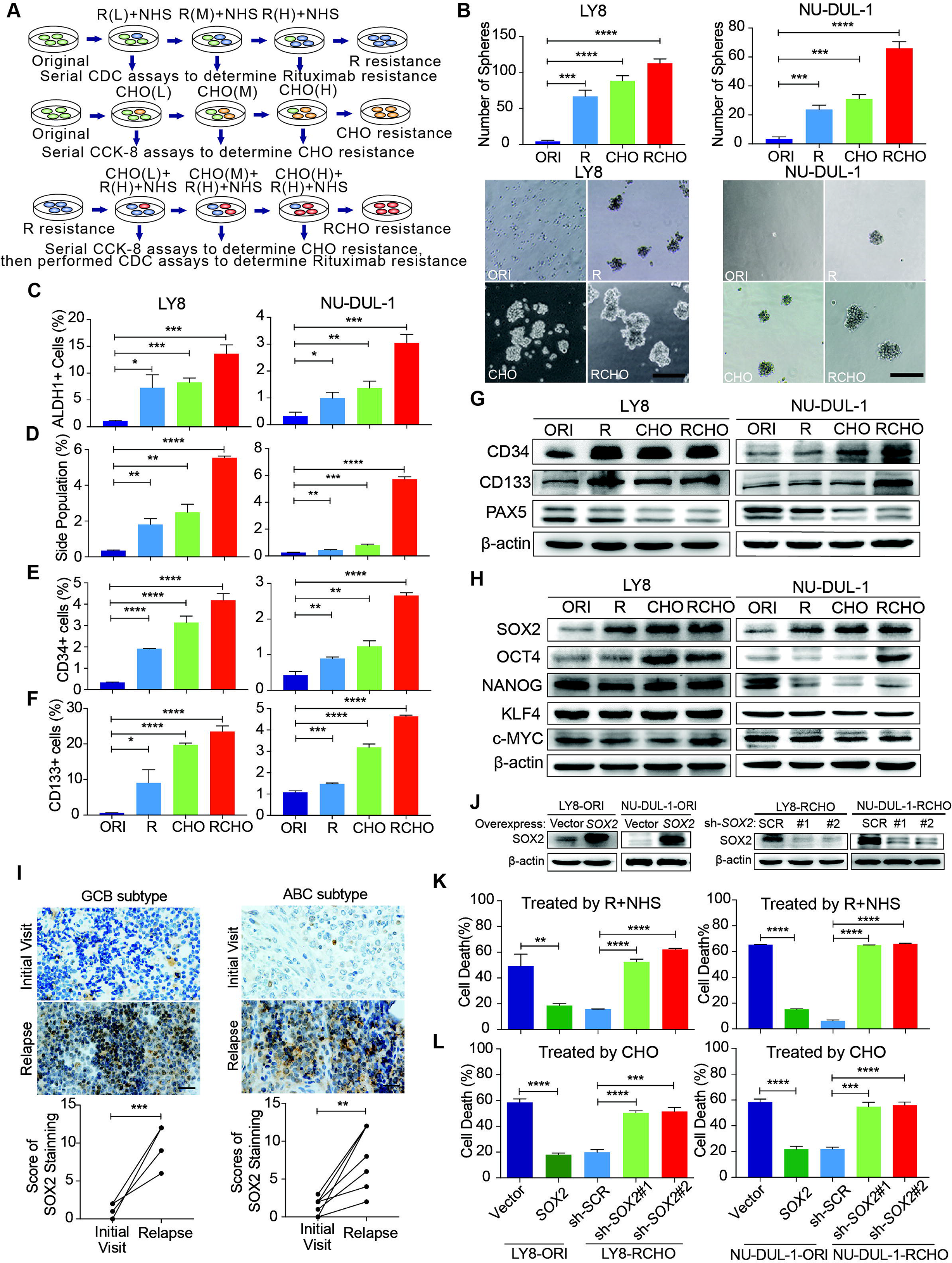
Resistant DLBCL cells potentiated stem-like features and their resistance was regulated by the SOX2 level. (**A**) A schematic diagram of the generation of resistant DLBCL cells by escalating drug concentration. L, M, and H represents the low, middle, and high dosages, respectively. (**B**-**G**) The proportion of TPCs in resistant DLBCL cells increased. The resistant cells exhibited enhanced sphere-forming capacity (B) and a greater percentage of ALDH1^+^ (C), side population fraction (D), CD34^+^ (E) and CD133^+^ cells (F) than original cells. Moreover, immunoblotting assay further revealed that CD34 and CD133 expressions increased, while PAX5 expression reduced in resistant cells (G). Scale bar: 100 μm. (**H**) Among the detected stemness-associated transcription factors, only SOX2 expression was gradually elevated along with the tendency toward resistance. (**I**) SOX2 expression significantly increased in the relapsed GCB (up, 6 pairs) and ABC (bottom, 6 pairs) subtype clinical tissues *vs* the paired tissues from the initial visit. Left: a representative image; Right: the quantitative score for SOX2 staining. Scale bar: 20 µm. (**J**) The SOX2 levels were confirmed by Immunoblotting assay after ectopic expressing SOX2 in the original cells and reducing SOX2 levels in RCHO-resistant cells. (**K**) CDC assays: ectopic SOX2 expression reduced the susceptibility of original cells to R-mediated CDC, whereas SOX2 silencing exhibited the opposite effect in RCHO-resistant cells. (**L**) CytoTox-Glo cytotoxicity assays: ectopic SOX2 expression of reduced the cytotoxic effect of CHO in original cells, whereas SOX2 silencing retrieved sensitivity to CHO in RCHO-resistant cells. The data are presented as mean ± SD; n=3. \**P*<0.05, \*\**P*<0.01, \*\*\**P*<0.001, and \*\*\*\**P*<0.0001.

**Table 1.**
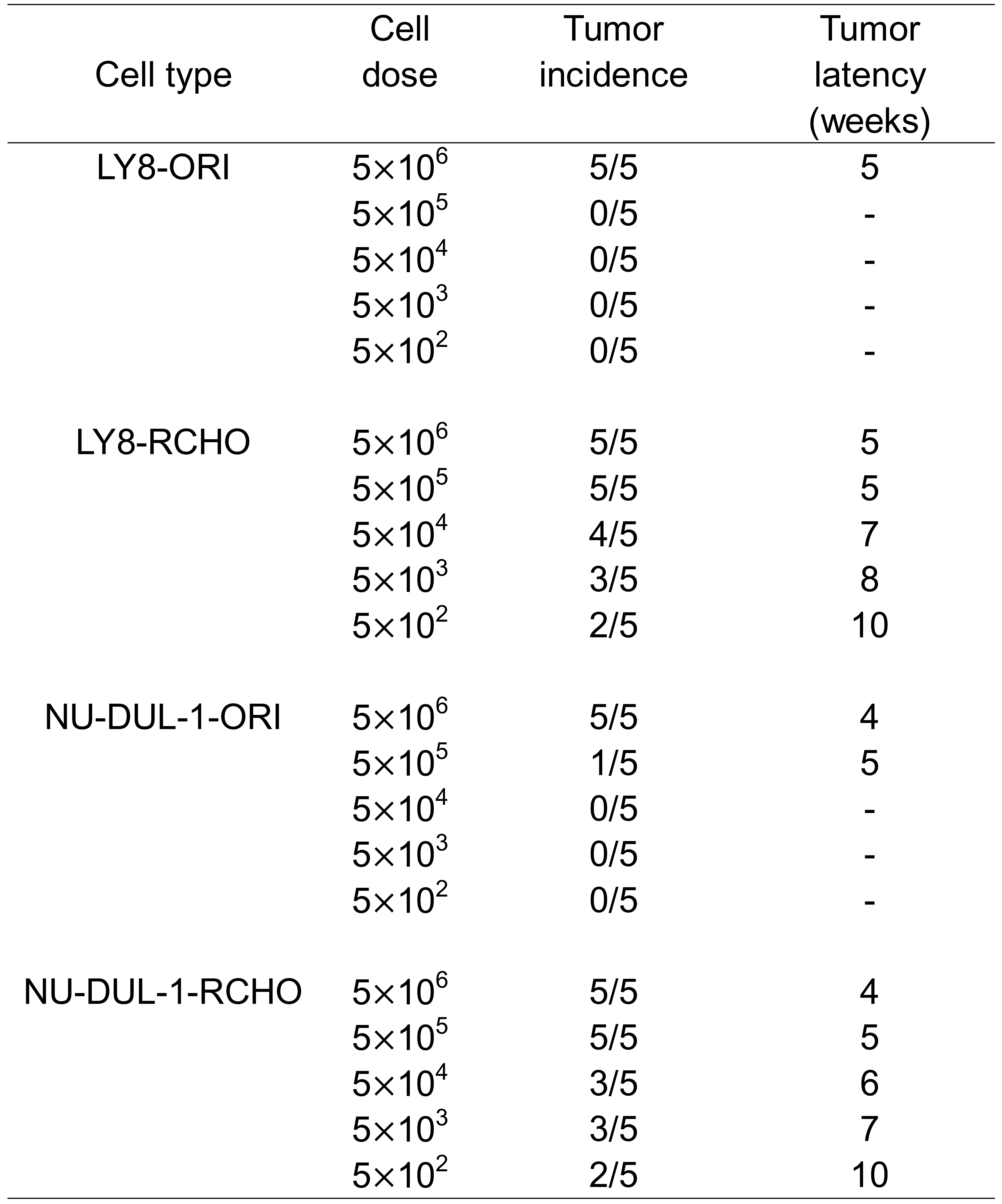
RCHO-resistant DLBCL cells display higher tumor-initiating capability than original DLBCL cells.

To examine the molecular regulation of stemness in the resistant cells, we detected the expression of core stemness-associated transcription factors, SOX2, OCT4, NANOG, KLF4 and c-Myc. As shown in Fig. 1H, only SOX2 level gradually increased along with the tendency toward resistance. Moreover, in clinical specimens from DLBCL patients, we observed that SOX2 staining in both GCB and ABC subtypes was markedly increased in relapsed tissues compared with their paired tissues from the initial visit (Fig. 1I and Fig. S3). Ectopic SOX2 expression (Fig. 1J) considerably reduced cell death induced by R (Fig. 1K) or CHO (Fig. 1L) in original LY8 or NU-DUL-1 cells; in contrast, Reduction of SOX2 expression (Fig. 1J) induced the opposite effect (Fig. 1K and L) in RCHO-resistant LY8 or NU-DUL-1 cells. In addition, although OCT4 level also increased in part of resistant DLBCL cells (Fig. 1H), the ectopic expression of OCT4 in original cells (Fig. S4A) failed to change its sensitivity to R (Fig. S4B) or CHO (Fig. S4C). These results revealed that TPCs strongly enriched in resistant DLBCL cells, and elevated SOX2 was responsible for the development of drug resistance to R and CHO.

### PI3K/AKT1 signaling phosphorylated and thus stabilized SOX2 against ubiquitination-mediated degradation

We next investigated the mechanism responsible for SOX2 upregulation in resistant DLBCL cells. Unexpectedly, the *SOX2* mRNA level significantly declined in resistant cells, particularly in resistant LY8 cells (Fig. 2A), indicating that transcriptional regulation did not contribute to SOX2 overexpression. Based on the RNA-seq data for the original and resistant cells, the interaction score from the KEGG pathway analysis showed that the PI3K/AKT signaling pathway and steroid biosynthesis were the most enriched pathways (Fig. 2B and Supplementary Data 1). Furthermore, gene set enrichment analysis (GSEA) demonstrated that the gene signatures for the PI3K/AKT signaling pathway were significantly activated in both RCHO-resistant cell lines (Fig. 2C and Supplementary Data 2). Experimentally, the result of flow cytometry also revealed that SOX2^+^ subpopulation exhibited higher AKT1 (S473) phosphorylation level than SOX2^−^ subpopulation in both RCHO-resistant LY8 and NU-DUL-1 cells (Fig. S5). Meanwhile, AKT1 (S473) phosphorylation was confirmed to remarkably increase in accordance with the resistance degree accompanied by the subsequently increased phosphorylation of its substrate PRAS40, thus enhancing SOX2 (T118) phosphorylation and subsequently suppressing SOX2 (K119) methylation in both cell lines. In addition, expression of the PI3K subunit p110α/δ and p85 in resistant LY8 cells and p110γ in resistant NU-DUL-1 cells were also elevated (Fig. 2D). Consistently, for both GCB and ABC subtypes, AKT1 (S473) phosphorylation was also significantly increased in the relapsed biopsy tissues versus the paired patient tissues from the initial visit (Fig. 2E and Fig. S6). The switch of SOX2 phosphorylation-methylation, which is regulated by PI3K/AKT1, was confirmed to regulate SOX2 stabilization via the ubiquitin system (*13*). Consistently, ubiquitinated-SOX2 was markedly reduced in resistant cells (Fig. 2F). Further, ectopic expression of a constitutively-active form of myristoylated AKT1 in original LY8 and NU-DUL-1 cells, which was determined by increased levels of p-AKT1 (S473) and p-PRAS40 (T246), potently elevated SOX2 level, thus inducing the strong resistance to R and CHO treatment (Fig. S7). Therefore, the elevated SOX2 expression in resistant TPCs of GCB and ABC subtypes may result directly from the AKT1-regulated phosphorylation-methylation switch, further inhibiting SOX2 ubiquitination and degradation.

**Fig. 2.**
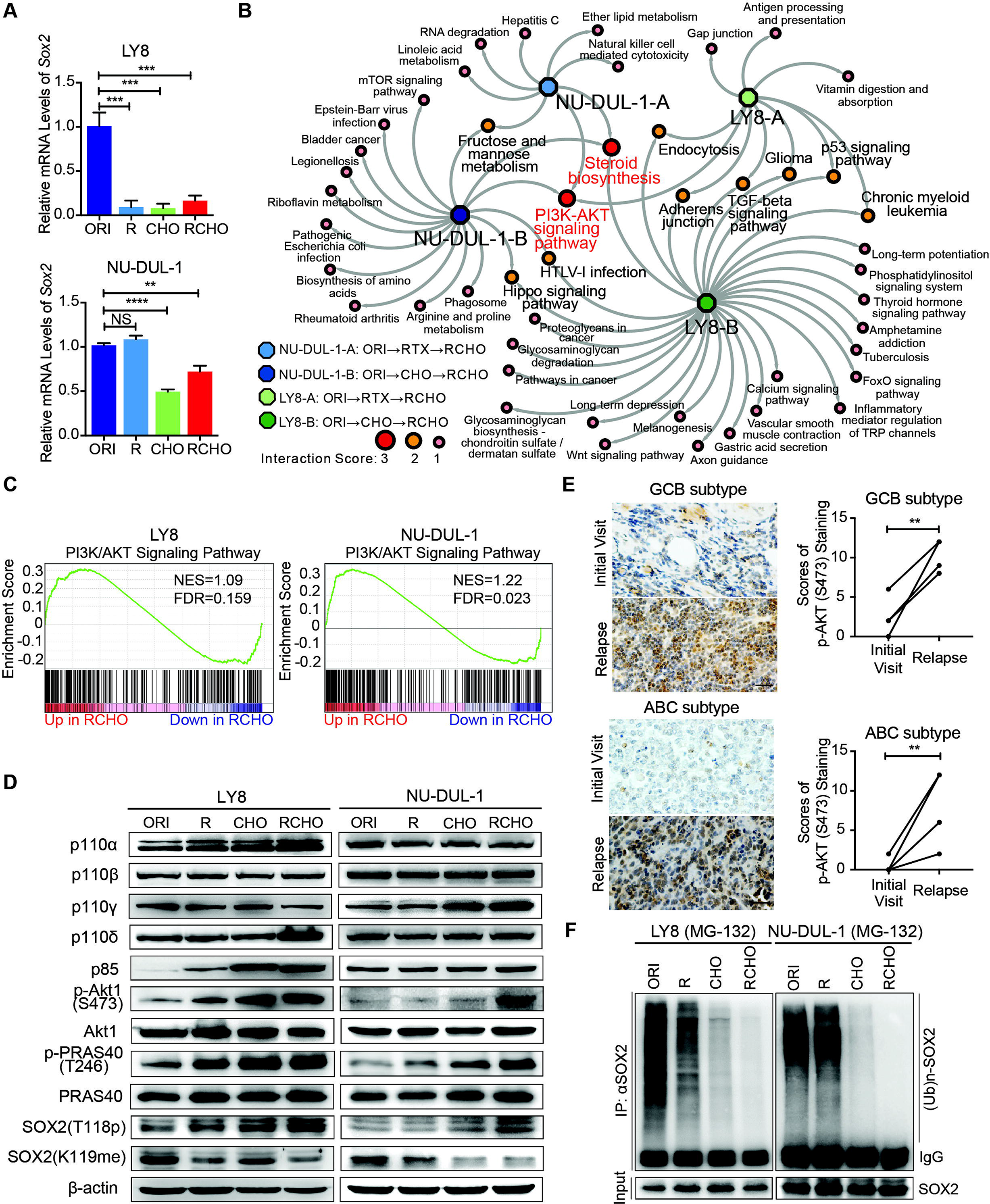
PI3K-AKT1 phosphorylates and further stabilizes SOX2 against ubiquitination-mediated degradation in resistant DLBCL cells. (**A**) The *SOX2* mRNA levels were significantly reduced in almost resistant DLBCL cells, as determined by qRT-PCR. (**B**) PI3K/AKT signaling together with steroid biosynthesis exhibited the highest interaction score in the network generated by Cytocape software using the RNA-seq data. (**C**) GSEA for the PI3K/AKT signature in RCHO-resistant LY8 (left) and NU-DUL-1 (right) cells. FDR<0.25 was considered significant. (**D**) PI3K/AKT1 signaling was activated, thus promoting the phosphorylation and reducing the methylation of SOX2. (**E**) PI3K/AKT1 signaling was markedly activated in relapsed GCB (up, 6 pairs) and ABC (bottom, 6 pairs) subtype clinical tissues *vs* the paired tissues from the initial visit. Left: a representative image; Right: the quantitative score for p-AKT1 (S473) staining. Scale bar: 20 μm. (**F**) Ubiquitination of SOX2 was reduced in resistant DLBCL cells after 10 μM MG-132 treatment for 8 hours. The data are presented as mean ± SD; n=3. \*\**P*<0.01, \*\*\**P*<0.001, and \*\*\*\**P*<0.0001.

### Multiple signaling events activated PI3K/AKT1 in resistant cells

An understanding of the key regulators upstream of the PI3K/AKT pathway is helpful for identification of potential drug targets to reduce SOX2 expression. Thus, we determined the up-regulated genes in the PI3K/AKT pathway (Fig. 2C and Supplementary Data 2) and classified them according to cell subtype (Fig. 3A). Due to the limited number (18) of overlapping genes in the two cell subtypes, we used all 124 genes to perform the KEGG pathway analysis (Fig. 3A). The top 10 pathways are listed in Fig. 3B and Supplementary Data 3, and PI3K/AKT was convincingly ranked as number 1. PI3Kα/β isoforms are universally expressed, whereas PI3Kγ/δ are exclusively limited to hematopoietic cells, in which PI3Kδ plays a critical role in B cell development and function. Assuming that a number of potential upstream signalings may contribute to PI3K/AKT activation in concert (Fig. 3B and Supplementary Data 3), we verified the reported activators, including BCR and other receptors to various integrins and cytokines/chemokines in lymphoma (*39–41*). Consistently, we found that the integrin-regulated focal adhesion pathway was ranked number 2 (Fig. 3B); however, downstream FAK was activated only in resistant LY8 cells (Fig. 3C). Next, we screened the transcription levels of all integrin subunits via qRT-PCR and found that only *ITGA1* and *ITGB5* were dramatically elevated in three resistant LY8 cell lines (Fig. 3D), which was confirmed by the protein levels (Fig. 3E). Reduction of *ITGA1* or *ITGB5* expression suppressed FAK and AKT1 phosphorylation, thus increasing SOX2 degradation (Fig. 3F). These results demonstrated that elevated expression of the integrin subunits α1 and β5 promoted SOX2 stabilization via FAK/AKT1 phosphorylation in resistant LY8 cells. In addition, the expression of CD79A, a constituent subunit of BCR signaling, and phosphorylation of downstream Lyn increased in both RCHO-resistant LY8 and NU-DUL-1 cells compared with the related original cells (Fig. 3G). These regulations eventually activated Syk phosphorylation in both RCHO-resistant cells (Fig. 3G). Further, the reduction of CD79A expression suppressed the BCR/Lyn/Syk/AKT1 signaling axis, thus inducing SOX2 instability (Fig. 3H). These results demonstrate that BCR signaling positively regulates the level of SOX2 protein (Fig. 3B). Moreover, we further investigated chemokine expression, which was suggested to activate the PI3K/AKT1 pathway (Fig. 3B). The GSEA results also showed that the chemokine signaling pathway was significantly enriched in RCHO-resistant LY8 cells but not in NU-DUL-1 cells (Fig. 3I, and Supplementary Data 4); the top 10 related genes are shown in Fig. 3J. We verified that CCR7 was ranked number 1 and found dramatically elevated *CCR7* mRNA levels in different resistant LY8 cells, especially RCHO-resistant cells (Fig. 3K). The CCR7 protein levels were consistently elevated, especially in CHO- and RCHO-resistant LY8 cells (Fig. 3L, left); in addition, CCR7 expression was elevated in resistant NU-DUL-1 cells, although to a lesser extent (Fig. 3L, right). Reduction of CCR7 expression impaired Src/AKT1 signaling, thus resulting in SOX2 degradation in both RCHO-resistant cell lines (Fig. 3M).

**Fig. 3.**
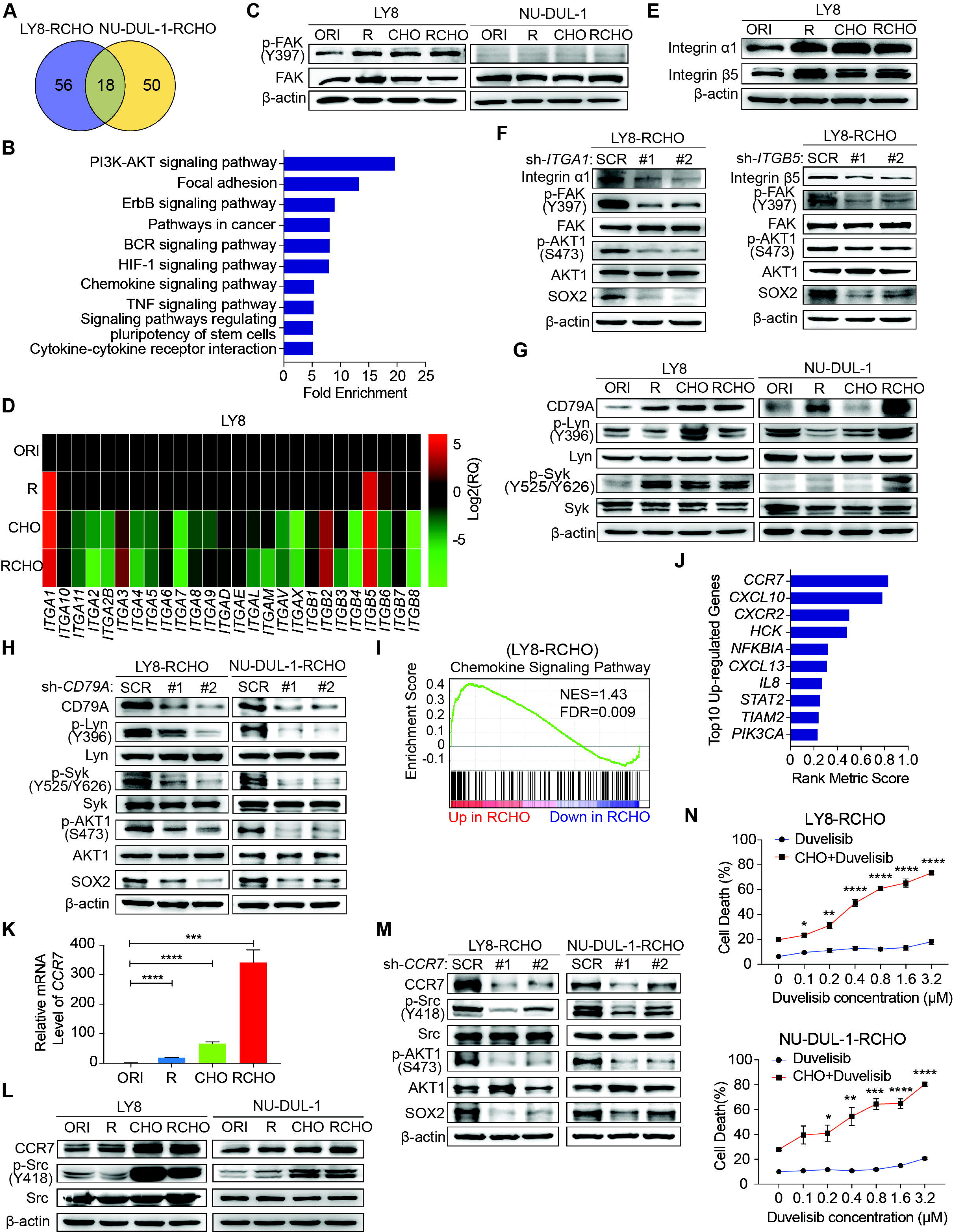
Multiple signaling events activated PI3K/AKT1 in RCHO-resistant cells. (**A**) Venn diagram illustrating the number of overlapping up-regulated genes in the PK3K/AKT1 pathway between RCHO-resistant LY8 and NU-DUL-1 cells. (**B**) The enriched top 10 signaling pathways based on the total 124 up-regulated genes in (A). (**C**) FAK was activated in resistant LY8 but not NU-DUL-1 cells. (**D**) The mRNA levels of the integrin subunits *ITGA1* and *ITGB5* were significantly up-regulated in resistant LY8 cells, as detected by qRT-PCR. (**E**) The integrin α1 and β5 subunits were up-regulated in resistant LY8 cells. (**F**) Knockdown of *ITGA1* or *ITGB5* suppressed the FAK-AKT1 signaling axis, leading to SOX2 degradation. (**G**) Syk was activated in resistant DLBCL cells by BCR-Lyn signaling pathways. (**H**) Silencing of the BCR component *CD79A* suppressed the BCR-Lyn-Syk-AKT1 signaling axis, leading to SOX2 degradation. (**I**) GSEA enriched chemokine signature in RCHO-resistant LY8 cells. FDR<0.25 was considered significant. (**J**) The top 10 up-regulated genes in the chemokine signaling pathway. (**K**) The *CCR7* mRNA level was significantly elevated in resistant LY8 cells, as determined by qRT-PCR. (**L**) CCR7 expression was up-regulated in CHO- and RCHO-resistant LY8 and NU-DUL-1 cells, thus enhancing downstream Src (Y418) phosphorylation. (**M**) Silencing of *CCR7* attenuated the CCR7-Src-AKT1 signaling axis, leading to SOX2 degradation. (**N**) CytoTox-Glo cytotoxicity assays: addition of the PI3K inhibitor duvelisib to CHO significantly reversed CHO resistance; however, duvelisib alone showed a negligible effect on direct induction of cell death. The data are presented as mean ± SD; n=3. \**P*<0.05, \*\**P*<0.01, \*\*\**P*<0.001, and \*\*\*\**P*<0.0001.

Next, we functionally tested the effect of inhibitors of the above signaling molecules on reversing resistance to chemotherapy (CHO) in resistant cells. Treatment with the PI3K inhibitor duvelisib alone induced marginal, if any, cytotoxicity; however, additive treatment with duvelisib and CHO dramatically enhanced the cytotoxic effect of CHO in both RCHO-resistant cell lines in a dose-dependent manner (Fig. 3N). Similarly, treatment with a FAK, Syk, or Src inhibitor alone induced negligible cytotoxicity; however, combination treatment with CHO significantly enhanced the cytotoxic effect of CHO, but to a much smaller degree than the duvelisib combination (Fig. S8). The sensitizing effect of duvelisib resulted most likely from the fact that duvelisib in advance reduced the proportion of TPCs in RCHO-resistant cells determined by the sphere formation assay and ALDH1^+^ cell measurement (Fig. S9). In addition, copanlisib, the first approved PI3K inhibitor with potent p110α/δ activities (*42*), displayed the similar effect with duvelisib that copanlisib also significantly sensitized the RCHO-resistant cells to CHO treatment due to AKT1 inhibition and resultant SOX2 degradation (Fig. S10). To summarize, the elevated PI3K/AKT1 signaling in resistant LY8 and NU-DUL-1 cells was induced by multiple factors, including integrins, BCR and chemokine signaling, and inhibition of PI3K/AKT1 but not of either upstream pathways effectively suppressed resistant cell survival during chemotherapy, perhaps by reducing TPC proportion.

### SOX2-regulated CD20 expression involved in the resistance development to R-induced CDC

We further investigated the mechanisms for the development of resistance to R-induced CDC. In both LY8 and NU-DUL-1 R- and RCHO-resistant cells, CD20 membrane expression was dramatically reduced, while the expression of three mCRPs was slightly decreased (Fig. S11A). Further, total CD20 level detected by immunoblotting showed the similar result (Fig. S11B), indicating that the significantly reduced membrane level of CD20 most likely resulted from transcription suppression (Fig. S11C).

The critical role of SOX2 in resistance to rituximab-mediated CDC in DLBCL has been previously demonstrated (Fig. 1J), and thus, we further examined the underlying mechanism. In both the original LY8 and NU-DUL-1 cells, ectopic SOX2 expression markedly reduced and SOX2 insufficiency dramatically increased CD20 expression (Fig. S11D). However, the expression of three mCRPs was not consistently associated with the resistance to rituximab-mediated CDC (Fig. S11D). These results suggest that CD20 expression regulated by SOX2 is the determining factor in DLBCL resistance to rituximab-mediated CDC compared with mCRPs.

We further determined the effect of PI3K/AKT inhibitors on rituximab-mediated CDC. Both PI3K and AKT suppression considerably reduced CD20 expression, while the expression levels of the three mCRPs were not significantly altered in either RCHO-resistant cell lines (Fig. S11E). Similarly, the reduced CD20 expression most likely resulted from suppressed transcription (Fig. S11F). AKT1 inhibition together with CD20 and SOX2 expression was confirmed by immunoblotting (Fig. S11G). These results also suggested that PI3K/AKT1 might regulate CD20 expression independently of SOX2. Subsequently, PI3K/AKT inhibition was found to impair rituximab-mediated CDC in RCHO-resistant cells (Fig. S11H). In addition, another PI3K inhibitor copanlisib also showed the similar effect with duvelisib that it conversely increased resistance to R treatment mostly by reducing CD20 level (Fig. S12). Therefore, in contrast with their effect on reversing resistance to chemotherapy, PI3K/AKT inhibitors might instead exacerbate resistance to rituximab-mediated CDC.

### GCB and ABC cells employed different SOX2-targeted signaling molecules to develop resistance

Given the effect of PI3K/AKT inhibition or SOX2 insufficiency on reversing resistance to R and/or CHO treatment, we analyzed up-regulated genes in the PI3K/AKT pathway (Fig. 2C and Supplementary Data 2) and SOX2-targeted genes (Molecular Signatures Database, MSigDB) to identify the critical targets of SOX2 in RCHO-resistant cells. Three genes, i.e., *CDK6*, *GSK3B*, and *SGK3*, were up-regulated in RCHO-resistant LY8 cells, while four genes, i.e., *FGFR1*, *FGFR2*, *KDR*, and *TNC*, were up-regulated in RCHO-resistant NU-DUL-1 cells (Fig. 4A and Supplementary Data 5). Considering that CDK4/6 (*43*) and FGFR (*44*) inhibitors are being tested in clinical trials for treatment of multi-type cancers, we subsequently investigated their expression and functions. CDK6 was highly expressed in resistant LY8 cells (Fig. 4B) and relapsed GCB but not in ABC subtype DLBCL tissues (Fig. 4C and Fig. S13). Consistently, FGFR1/2 were overexpressed in resistant NU-DUL-1 cells (Fig. 4D) and relapsed ABC but not in GCB subtype DLBCL tissues (Fig. 4E and F, and Fig. S14 and S15). Moreover, SOX2^+^ subpopulation expressed higher CDK6 or FGFR1/2 than SOX2^−^ subpopulation in RCHO-resistant LY8 or NU-DUL-1 cells, respectively (Fig. S16), further indicating the close association between SOX2 and CDK6 or FGFR1/2. PI3K inhibition with duvelisib suppressed AKT1 and PRAS40 phosphorylation and induced SOX2 degradation, thus reducing CDK6 expression in LY8 RCHO-resistant cells (Fig. 4G) or FGFR1/2 expression in NU-DUL-1 RCHO-resistant cells (Fig. 4H) in a time dependent manner. We observed that SOX2 and OCT4 levels reduced starting from 2 or 24 (Fig. S4D) hours, respectively, after duvelisib treatment in both RCHO-resistant cells, indicating the different regulation of SOX2 and OCT4 in the resistant cells. Duvelisib effect on LY8 RCHO-resistant cells was more transient than that in NU-DUL-1 RCHO-resistant cells (Fig. 4G and H), which resulted probably from the diverse metabolic features in different cell types. Duvelisib also displayed dose-effect relationship with above molecules (Fig. 4I and J). Similar to the effect of PI3K inhibition, CDK6 inhibition with abemaciclib in RCHO-resistant LY8 cells or FGFR1/2 inhibition with AZD454 in RCHO-resistant NU-DUL-1 cells considerably enhanced the cytotoxicity of CHO, while these inhibitors alone displayed a negligible cytotoxic effect on the resistant cells (Fig. 4K and L). However, CDK6 inhibition in LY8-RCHO or FGFR1/2 inhibition in NU-DUL-1-RCHO-resistant cells failed to reverse resistance to R-mediated CDC (Fig. 4M and N). These results indicated that inhibition of SOX2 targets (CDK6 in LY8-RCHO-resistant cells and FGFR1/2 in NU-DUL-1-RCHO-resistant cells) could at least partly reverse resistance to chemotherapy but not to R-mediated CDC.

**Fig. 4.**
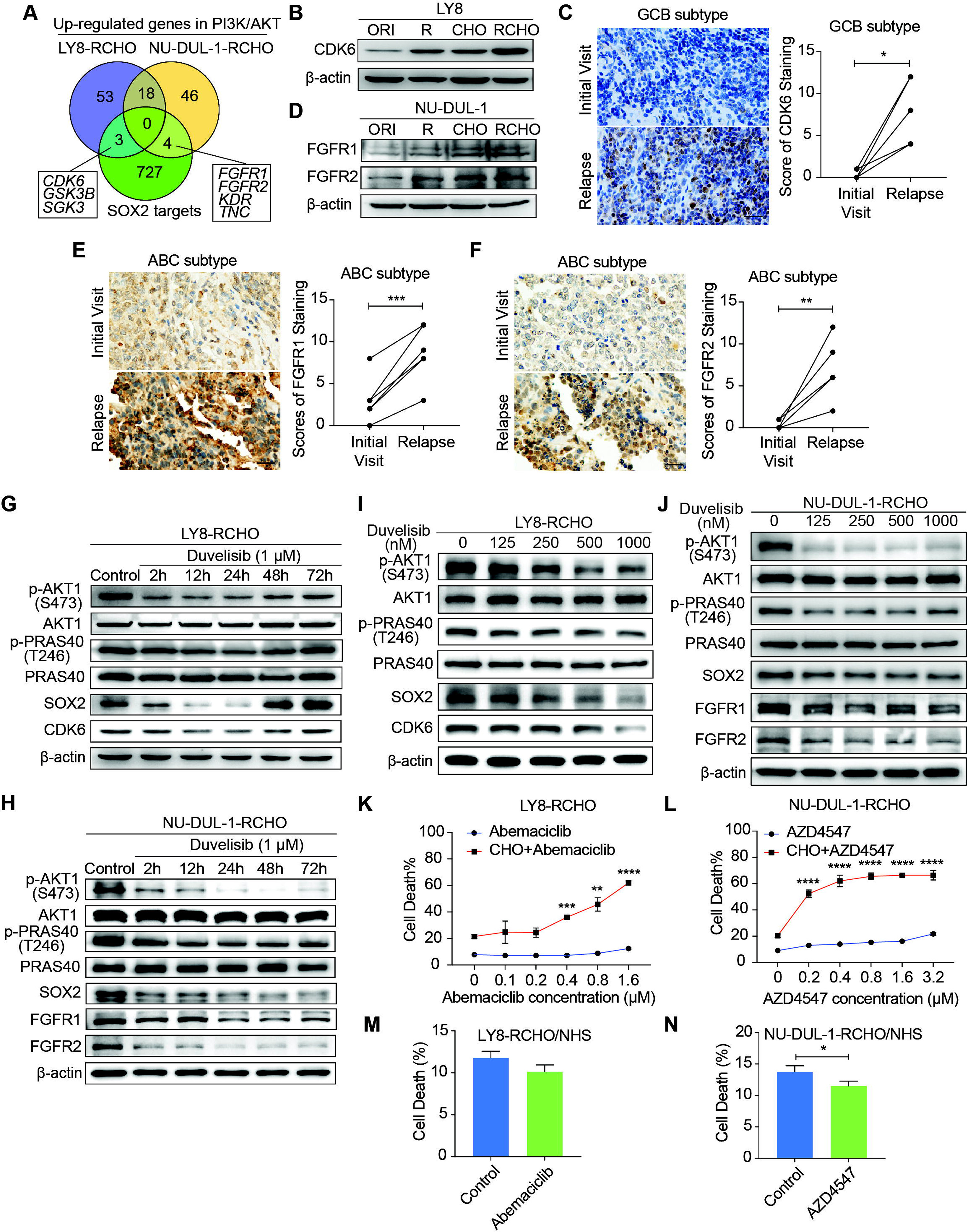
Inhibition of CDK6 or FGFR1/2 restored sensitivity to CHO treatment. (**A**) Venn diagram illustrating the number of overlapping up-regulated genes in the PK3K/AKT1 pathway and SOX2 targets. (**B**) CDK6 was overexpressed in resistant LY8 cells. (**C**) CDK6 was up-regulated in relapsed tissues compared with the paired tissues from the initial visit in 6 patients with the GCB subtype. (**D**) FGFR1 and FGFR2 were overexpressed in resistant NU-DUL-1 cells. (**E**-**F**) FGFR1 (E) and FGFR2 (F) was up-regulated in relapsed tissues compared with the paired tissue from the initial visit in 6 patients with the ABC subtype. Left: a representative image (C, E and F); Right: the quantitative scores for CDK6 (C), FGFR1 (E) and FGFR2 (F) staining. Scale bar: 20 μm. **(G**-**J)** Duvelisib reduced the expression of SOX2 and CDK6 (G and I) or FGFR1/2 (H and J) by suppressing AKT1 activity in RCHO-resistant LY8 (G and I) or NU-DUL-1 (H and J) cells, respectively, in a time- (G-H) and dose- (I-J) dependent manner. (**K**-**L**) CytoTox-Glo cytotoxicity assay: inhibition of CDK6 by abemaciclib in RCHO-resistant LY8 cells (K) or inhibition of FGFR1/2 by AZD4547 in RCHO-resistant NU-DUL-1 cells (L) restored sensitivity to CHO treatment. Abemaciclib or AZD4547 alone exhibited a negligible effect on induction of cell death. (**M**-**N**) CDC assay: abemaciclib (M) failed to reverse and AZD4547 (N) exacerbated resistance to R-mediated CDC in the indicated RCHO-resistant DLBCL cells. The data are presented as mean ± SD; n=3. \**P*<0.05, \*\**P*<0.01, \*\*\**P*<0.001, and \*\*\*\**P*<0.0001.

### PI3K/AKT Inhibition promoted TPCs differentiation by reducing SOX2

We next investigated the role of PI3K/AKT/SOX2 axis in stemness maintenance. The expression levels of CD34, CD133 were clearly reduced after PI3K inhibition in both RCHO-resistant cell lines (Fig 5A), whereas CDK6 inhibition had no significant effect on the expression of the above molecules (Fig 5A). However, the expression levels of the above molecules also decreased after FGFR1/2 inhibition but to a smaller extent than those after PI3K inhibition (Fig 5A). In addition, PI3K inhibition significantly reduced the proportion of TPCs (marked as ALDH1^+^ cells) in both RCHO-resistant cell lines (Fig. 5B and C); whereas CDK6 inhibition failed to change the proportion of TPCs in LY8 RCHO-resistant cells (Fig. 5B). However, FGFR1/2 inhibition also significantly reduced the proportion of TPCs in NU-DUL-1 RCHO-resistant cells but to a much smaller degree than that with PI3K inhibition (Fig. 5C), in agreement with a previous report showing that PI3K/AKT1 may be activated by FGFR signaling (*45*). Notably, FGFR1/2 were up-regulated and participated in PI3K/AKT1 signaling activation (Supplementary Data 3); thus, their inhibition reduced AKT1 phosphorylation and SOX2 stabilization (Fig. 5A). These results revealed that PI3K/AKT1 inhibition directly by duvelisib or indirectly by AZD4547 promoted TPCs differentiation.

**Fig. 5.**
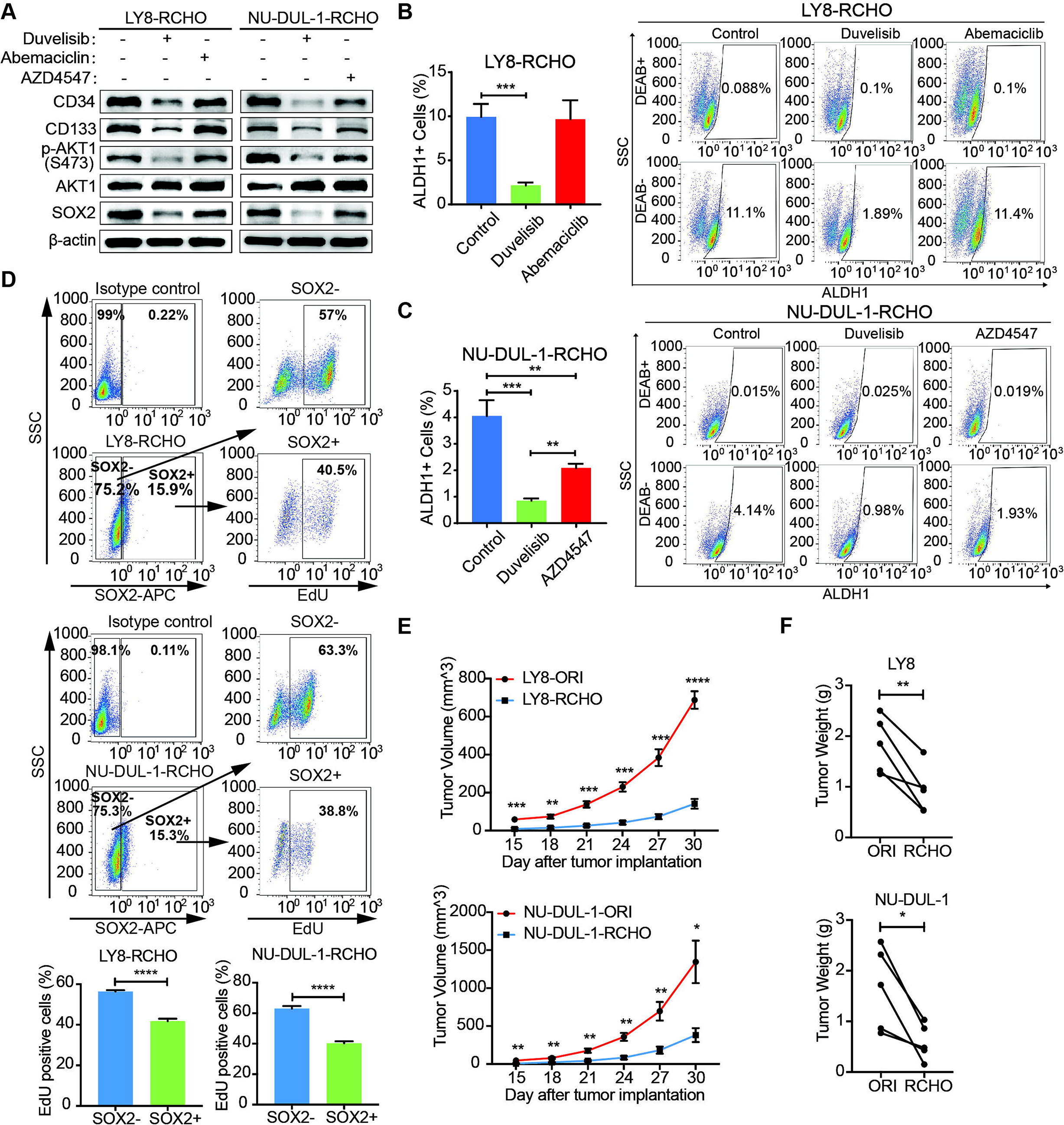
PI3K/AKT inhibition promoted differentiation of resistant DLBCL cells by reducing SOX2 expression. (**A**) Duvelisib clearly reduced the levels of CD34, CD133 in RCHO-resistant cells; abemaciclib exhibited no effect on their levels; while AZD4547 also reduced their levels, but to a greatly reduced degree compared with duvelisib. (**B**-**C**) Duvelisib dramatically reduced the ALDH1^+^ subpopulation in both RCHO-resistant LY8 (B) and NU-DUL-1 (C) cells. Abemaciclib had no effect on the ALDH1^+^ subpopulation in RCHO-resistant LY-8 cells (B), while AZD4547 significantly reduced this subpopulation in RCHO-resistant NU-DUL-1 cells, but to a greatly reduced extent compared with duvelisib (C). (**D**) SOX2^+^ subpopulation showed less EdU positive cells than SOX2^−^ population in RCHO-resistant LY8 (up) and NU-DUL-1 (down) cells detected by flow cytometry. Representative images (up) and quantitative results (bottom). The data are presented as mean ± SD; n=3. \*\**P*<0.01, \*\*\**P*<0.001, and \*\*\*\**P*<0.0001. (**E**-**F**) Both RCHO-resistant LY8 and NU-DUL-1 cells grew more slowly than the related original cells in the *in vivo* serial-transplantation experiment with 5×10^6^ tumor cells injection, in which both RCHO-resistant and original cells induced 100% tumor incidence (Table 1). Tumor growth rate (E) and paired tumor weight (F) at end-point of experiment. The data are presented as mean ± SEM; n=5. \**P*<0.05, \*\**P*<0.01, \*\*\**P*<0.001, and \*\*\*\**P*<0.0001.

Differentiated tumor cells grow faster than TPCs. Indeed, SOX2^+^ subpopulation showed less EdU positive cells than SOX2^−^ subpopulation in RCHO-resistant LY8 and NU-DUL-1 cells (Fig. 5D). Further, both RCHO-resistant cells grew more slowly than their related original cells in the *in vivo* serial-transplantation experiment with 5×10^6^ cells injection (Fig. 5E and F), in which both RCHO-resistant and original cells induced 100% tumor incidence (Table 1). PI3K/AKT inhibition effectively converted TPCs to differentiated cells, while CDK6 inhibition might directly suppress TPC growth through cell cycle targeting, and FGFR1/2 might have a dual effect on TPCs. All of these interventions eventually enhanced the TPC susceptibility to chemotherapy with varying efficacy.

### Pro-differentiation therapy against TPCs by combining a PI3K inhibitor with R-CHOP suppressed tumor growth of RCHO-resistant cells

RCHO-resistant LY8 and NU-DUL-1 cells transduced with luciferase-expressing plasmid were separately implanted into immuno-deficient mice, and various therapeutic regimens were administered. In mice bearing RCHO-resistant LY8 cells, R-CHOP treatment had no therapeutic effect on tumor growth compared with the saline control (Fig. 6A and B), although R-CHOP significantly prolonged the survival rate (Fig. 6C). These results suggested that the *in vitro* resistance of LY8 cells to RCHO was recapitulated in mice to a high degree. Neither duvelisib nor abemaciclib alone suppressed tumor growth or prolonged the survival rate (Fig. 6A-C); however, duvelisib accelerated tumor growth compared with saline, and although this difference was not statistically significant (Fig. 6A and B), it led to a significantly shortened survival rate (Fig. 6C). In contrast, when combined with R-CHOP, duvelisib or abemaciclib dramatically suppressed tumor growth and prolonged survival compared with R-CHOP or inhibitor alone (Fig. 6A-C). More importantly, duvelisib combination therapy effectively suppressed tumorigenesis, and all the treated mice survived until the experimental end-point at day 120 compared with two mice receiving abemaciclib combination therapy that died on days 62 and 78 (Fig. 6A-C). Similar but not identical results were obtained in mice bearing RCHO-resistant NU-DUL-1 cells. R-CHOP significantly suppressed tumor growth and prolonged survival (Fig. 6D-F), likely due to the weaker stemness and reduced TPC proportions, resulting in weaker resistance than RCHO-resistant LY8 cells (Fig. 1 B-F). Duvelisib alone also had no therapeutic effect on tumor growth and survival, whereas AZD4547 alone significantly prolonged survival (Fig. 6D-F). Importantly, when combined with R-CHOP, both duvelisib and AZD4547 effectively suppressed tumor growth and prolonged survival up to the experimental end-point of day 120 (Fig. 6D-F). These different tumor-suppressing effects of duvelisib, abemaciclib and AZD4547 further support their potentially distinct resistance-reversing mechanisms; i.e., duvelisib converted TPCs to differentiated cells, abemaciclib inhibited TPC growth, and AZD4547 exhibited dual effects on TPCs. SOX2 staining of the implanted tumor tissues with different treatments supported this finding. Duvelisib or AZD4547, but not R-CHOP or abemaciclib, dramatically reduced SOX2 expression (Fig. 6G and H). Therefore, the success of this combination therapy most likely resulted from TPCs conversion to differentiated cells via PI3K/AKT1/SOX2 axis inhibition.

**Fig. 6.**
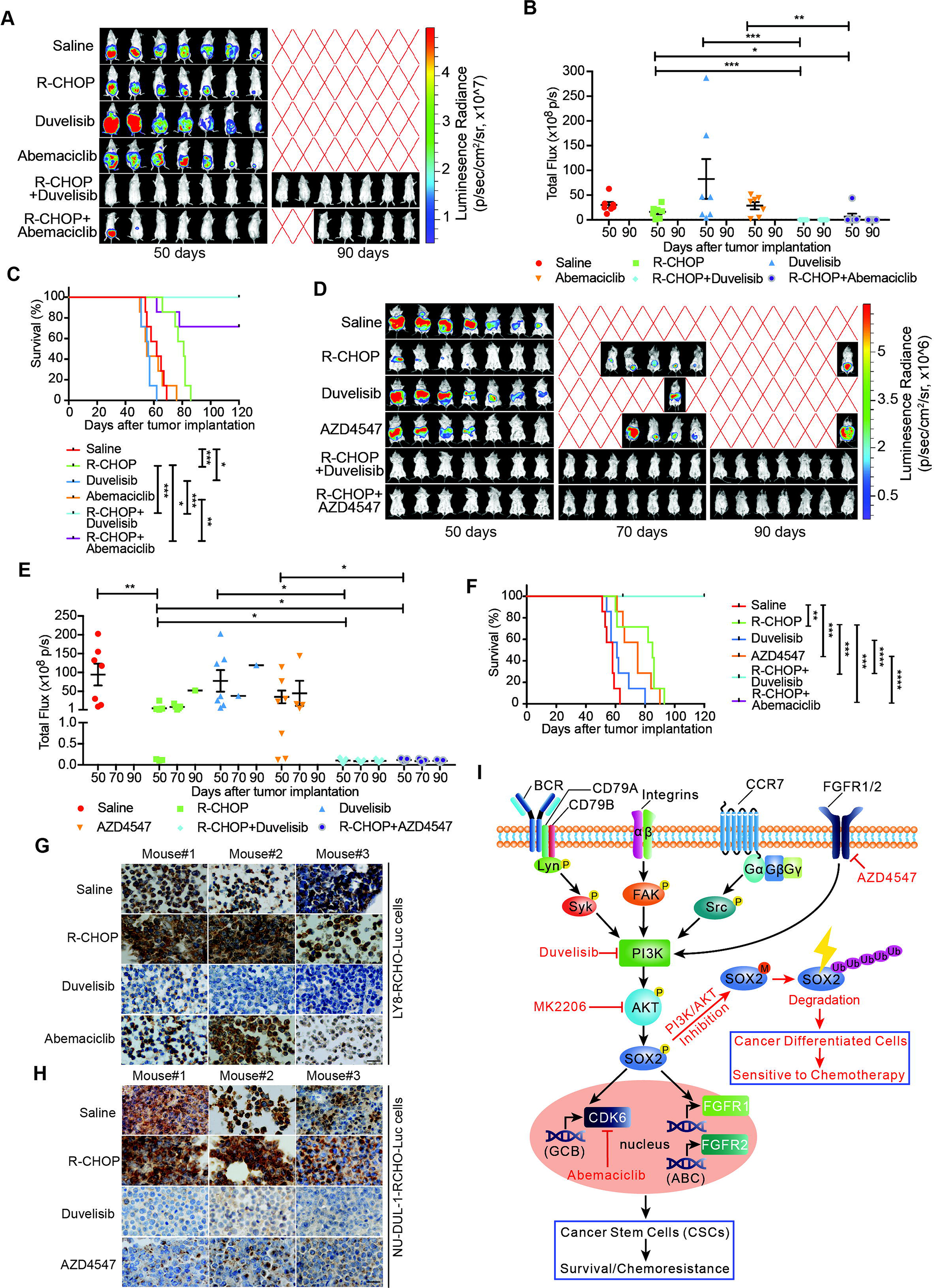
The combination of PI3K-AKT1 or FGFR1/2 inhibitors but not of CDK6 inhibitor with R-CHOP suppressed the tumor growth of RCHO-resistant cells. (**A**-**C**) Addition of abemaciclib and addition of duvelisib to R-CHO respectively, suppressed tumor growth of RCHO-resistant LY8 cells. (**D**-**F**) Addition of AZD4547 or duvelisib to R-CHOP effectively suppressed tumor growth of RCHO-resistant NU-DUL-1 cells. In vivo bioluminescence imaging (A and D). The tumor mass is represented by the quantified total photon flux (B and E). Kaplan–Meier survival curves (C and F). The red “X” in (A) and (D) represents mouse death. The data are presented as mean ± SEM (n=7). \**P*<0.05, \*\**P*<0.01, \*\*\**P*<0.001, and \*\*\*\**P*<0.0001. (**G**-**H**) SOX2 staining in LY8-RCHO-Luc (G) or NU-DUL-1-RCHO-Luc (H) cell-derived tumor tissues treated with the indicated drugs. Duvelisib and AZD4547, but not R-CHOP and abemaciclib, dramatically reduced the SOX2 expression level compared with the saline control. Scale bar, 20 μm. (**I**) Schematic showing the mechanisms by which pro-differentiation therapy against TPCs reverses drug resistance to R-CHOP in DLBCL.

## Discussion

Along with cancer progression, the differentiated phenotypes are gradually lost, and stem-like features are subsequently acquired by TPCs, resulting in metastasis and resistance to available therapies (*1, 2, 46–48*). However, the current attempts to reverse drug resistance by eradicating TPCs directly or slowing TPC growth indirectly have been far from successful, include direct inhibition via TPC-dependent signaling (Wnt/β-catenin, Hedgehog, Notch, FAK, PTEN, Nanog, and JAK/STAT, among others) inhibitors, tumor microenvironment modulators, direct eradication via CD44-targeting or CD133-targeting immunotherapy, and differentiation therapy (*7, 49*).

Herein, we provide a distinct strategy for targeting TPCs, termed pro-differentiation therapy (PDT). In RCHO-resistant DLBCL cells, BCR-mediated Syk, integrins, CCR7 and FGFR1/2 signaling could activate PI3K/AKT1, which subsequently stabilized SOX2 by enhancing SOX2 phosphorylation and reducing SOX2 methylation. Eventually, the up-regulated SOX2 increased the survival and proportion of TPCs through CDK6 or FGFR1/2, depending on the resistant DLBCL cell subtype, thus inducing resistance. When combined with R-CHOP, PI3K/AKT1 signaling inhibitors effectively reversed drug resistance by accelerating SOX2 degradation, subsequently promoting TPC differentiation and the resultant hypersensitivity to chemotherapy (Fig. 6I). Interestingly, using oncogenic dedifferentiation by machine learning in almost 12,000 samples of 33 tumor types, SOX2 has very recently been identified as a master stemness-associated transcription factor (*48*).

Given the critical role of TPCs in metastasis and drug resistance, we propose a novel strategy of PDT for coping with TPCs; i.e., identify the determinant signaling pathway for stemness maintenance and then guide TPCs to differentiate by interference with this pathway. The differentiated cells will eventually become sensitive to conventional therapies such as chemotherapy. In this case, a PI3K/AKT inhibitor regimen combined with R-CHOP merits assessment in clinical trials for resistant DLBCL patients.

## Acknowledgments

This work was supported by grants to W.H. from the National Natural Science Foundation of China (81372258) and from the Shanghai Science and Technology Committee (16JC1405500) and a grant to J.C. from the National Natural Science Foundation of China (81700190). We thank Ping Zhang and Yuhu Xin for technical support. Finally, we thank the DLBCL patients and their family members for supporting our research. RNA-seq data reported in this study are accessible in the NCBI’s GEO database through the GEO Series accession number GSE112989 and GSE113001.

## Authorship Contributions

W.H. conceived the project. W.H. and J.C. designed the experiments. J.C., X.G., W.Z., Y.D., Q.W., L.L., Y.S., P.Z., Y.Z., L.Z., P.D., and X.Z. performed most of the experiments. J.C. and H.G. performed the pathological experiments. J.C., X.G., and Y.D. analyzed the pathological data. L.F. and J.W. prepared the antibodies against phospho-SOX2 and me-SOX2. Q.Z., K.X., Q.L., and J.W. contributed helpful discussions about the project. J.C. and W.H analyzed the data, prepared the figures and wrote the paper.

## References

1. T. Shibue, R. A. Weinberg, EMT, CSCs, and drug resistance: the mechanistic link and clinical implications. Nature reviews. Clinical oncology 14, 611–629 (2017).

2. M. Dean, T. Fojo, S. Bates, Tumour stem cells and drug resistance. Nat Rev Cancer 5, 275–284 (2005).

3. C. E. Meacham, S. J. Morrison, Tumour heterogeneity and cancer cell plasticity. Nature 501, 328–337 (2013).

4. P. N. Kelly, A. Dakic, J. M. Adams, S. L. Nutt, A. Strasser, Tumor growth need not be driven by rare cancer stem cells. Science 317, 337 (2007).

5. E. Quintana et al., Efficient tumour formation by single human melanoma cells. Nature 456, 593–598 (2008).

6. J. Kaiser, The cancer stem cell gamble. Science 347, 226–229 (2015).

7. S. B. Nandy, R. Lakshmanaswamy, Cancer Stem Cells and Metastasis. Prog Mol Biol Transl Sci 151, 137–176 (2017).

8. A. A. Avilion et al., Multipotent cell lineages in early mouse development depend on SOX2 function. Genes & development 17, 126–140 (2003).

9. K. Takahashi, S. Yamanaka, Induction of pluripotent stem cells from mouse embryonic and adult fibroblast cultures by defined factors. Cell 126, 663–676 (2006).

10. E. L. Wuebben, A. Rizzino, The dark side of SOX2: cancer - a comprehensive overview. Oncotarget 8, 44917–44943 (2017).

11. J. R. Testa, A. Bellacosa, AKT plays a central role in tumorigenesis. Proc Natl Acad Sci U S A 98, 10983–10985 (2001).

12. P. Xia, X. Y. Xu, PI3K/Akt/mTOR signaling pathway in cancer stem cells: from basic research to clinical application. American journal of cancer research 5, 1602–1609 (2015).

13. L. Fang et al., A methylation-phosphorylation switch determines Sox2 stability and function in ESC maintenance or differentiation. Molecular cell 55, 537–551 (2014).

14. R. L. Siegel, K. D. Miller, A. Jemal, Cancer statistics, 2016. CA: a cancer journal for clinicians 66, 7–30 (2016).

15. A. A. Alizadeh et al., Distinct types of diffuse large B-cell lymphoma identified by gene expression profiling. Nature 403, 503–511 (2000).

16. J. O. Armitage, My treatment approach to patients with diffuse large B-cell lymphoma. Mayo Clinic proceedings 87, 161–171 (2012).

17. J. W. Friedberg, Relapsed/refractory diffuse large B-cell lymphoma. Hematology. American Society of Hematology. Education Program 2011, 498–505 (2011).

18. R. Camicia, H. C. Winkler, P. O. Hassa, Novel drug targets for personalized precision medicine in relapsed/refractory diffuse large B-cell lymphoma: a comprehensive review. Mol Cancer 14, 207 (2015).

19. G. S. Nowakowski et al., Beyond RCHOP: A Blueprint for Diffuse Large B Cell Lymphoma Research. J Natl Cancer Inst 108, (2016).

20. W. Hu et al., Human CD59 inhibitor sensitizes rituximab-resistant lymphoma cells to complement-mediated cytolysis. Cancer Res 71, 2298–2307 (2011).

21. S. A. Maxwell, E. M. Cherry, K. J. Bayless, Akt, 14-3-3zeta, and vimentin mediate a drug-resistant invasive phenotype in diffuse large B-cell lymphoma. Leukemia & lymphoma 52, 849–864 (2011).

22. C. Ginestier et al., ALDH1 is a marker of normal and malignant human mammary stem cells and a predictor of poor clinical outcome. Cell stem cell 1, 555–567 (2007).

23. J. P. Chute et al., Inhibition of aldehyde dehydrogenase and retinoid signaling induces the expansion of human hematopoietic stem cells. Proc Natl Acad Sci U S A 103, 11707–11712 (2006).

24. J. Chen et al., CD59 Regulation by SOX2 Is Required for Epithelial Cancer Stem Cells to Evade Complement Surveillance. Stem Cell Reports 8, 140–151 (2017).

25. Z. Liu et al., Autism-like behaviours and germline transmission in transgenic monkeys overexpressing MeCP2. Nature 530, 98–102 (2016).

26. M. R. Smith, Rituximab (monoclonal anti-CD20 antibody): mechanisms of action and resistance. Oncogene 22, 7359–7368 (2003).

27. X. Zhou, W. Hu, X. Qin, The role of complement in the mechanism of action of rituximab for B-cell lymphoma: implications for therapy. Oncologist 13, 954–966 (2008).

28. G. Cartron et al., Therapeutic activity of humanized anti-CD20 monoclonal antibody and polymorphism in IgG Fc receptor FcgammaRIIIa gene. Blood 99, 754–758 (2002).

29. A. Russ et al., Blocking “don’t eat me” signal of CD47-SIRPalpha in hematological malignancies, an in-depth review. Blood Rev, (2018).

30. Z. Fishelson, Obstacles to cancer immunotherapy: expression of membrane complement regulatory proteins (mCRPs) in tumors. Molecular Immunology 40, 109–123 (2003).

31. J. A. Martinez-Climent, L. Fontan, R. D. Gascoyne, R. Siebert, F. Prosper, Lymphoma stem cells: enough evidence to support their existence? Haematologica 95, 293–302 (2010).

32. E. Gross, A. Quillet-Mary, L. Ysebaert, G. Laurent, J. J. Fournie, Cancer stem cells of differentiated B-cell malignancies: models and consequences. Cancers 3, 1566–1579 (2011).

33. I. Ma, A. L. Allan, The role of human aldehyde dehydrogenase in normal and cancer stem cells. Stem cell reviews 7, 292–306 (2011).

34. L. Patrawala et al., Side population is enriched in tumorigenic, stem-like cancer cells, whereas ABCG2+ and ABCG2- cancer cells are similarly tumorigenic. Cancer Res 65, 6207–6219 (2005).

35. T. Lapidot et al., A cell initiating human acute myeloid leukaemia after transplantation into SCID mice. Nature 367, 645–648 (1994).

36. N. Feller et al., Immunologic purging of autologous peripheral blood stem cell products based on CD34 and CD133 expression can be effectively and safely applied in half of the acute myeloid leukemia patients. Clinical cancer research : an official journal of the American Association for Cancer Research 11, 4793–4801 (2005).

37. S. Lombardi et al., Growth hormone is secreted by normal breast epithelium upon progesterone stimulation and increases proliferation of stem/progenitor cells. Stem Cell Reports 2, 780–793 (2014).

38. I. Mikkola, B. Heavey, M. Horcher, M. Busslinger, Reversion of B cell commitment upon loss of Pax5 expression. Science 297, 110–113 (2002).

39. C. A. Durand et al., Phosphoinositide 3-kinase p110 delta regulates natural antibody production, marginal zone and B-1 B cell function, and autoantibody responses. J Immunol 183, 5673–5684 (2009).

40. A. Bilancio et al., Key role of the p110delta isoform of PI3K in B-cell antigen and IL-4 receptor signaling: comparative analysis of genetic and pharmacologic interference with p110delta function in B cells. Blood 107, 642–650 (2006).

41. B. J. Lannutti et al., CAL-101, a p110delta selective phosphatidylinositol-3-kinase inhibitor for the treatment of B-cell malignancies, inhibits PI3K signaling and cellular viability. Blood 117, 591–594 (2011).

42. N. Liu et al., BAY 80-6946 is a highly selective intravenous PI3K inhibitor with potent p110alpha and p110delta activities in tumor cell lines and xenograft models. Mol Cancer Ther 12, 2319–2330 (2013).

43. B. O’Leary, R. S. Finn, N. C. Turner, Treating cancer with selective CDK4/6 inhibitors. Nature reviews. Clinical oncology 13, 417–430 (2016).

44. I. S. Babina, N. C. Turner, Advances and challenges in targeting FGFR signalling in cancer. Nat Rev Cancer 17, 318–332 (2017).

45. S. H. Ong et al., Stimulation of phosphatidylinositol 3-kinase by fibroblast growth factor receptors is mediated by coordinated recruitment of multiple docking proteins. Proc Natl Acad Sci U S A 98, 6074–6079 (2001).

46. R. Pardal, M. F. Clarke, S. J. Morrison, Applying the principles of stem-cell biology to cancer. Nat Rev Cancer 3, 895–902 (2003).

47. B. M. Boman, M. S. Wicha, Cancer stem cells: a step toward the cure. Journal of clinical oncology : official journal of the American Society of Clinical Oncology 26, 2795–2799 (2008).

48. T. M. Malta et al., Machine Learning Identifies Stemness Features Associated with Oncogenic Dedifferentiation. Cell 173, 338–354 e315 (2018).

49. S. Talukdar, L. Emdad, S. K. Das, D. Sarkar, P. B. Fisher, Evolving Strategies for Therapeutically Targeting Cancer Stem Cells. Adv Cancer Res 131, 159–191 (2016).

