## Supplemental Information for "PI3K/AKT inhibition in tumor propagating cells of DLBCL reverses R-CHOP resistance by destabilizing SOX2"

**Inventory of Supplementary Information Items**

**ONLINE EXPERIMENTAL PROCEDURES**

**Supplemental References**

**Supplemental Figures and Legends**

**Supplemental Table S1**

**Supplemental Table S2**

**Supplemental Table S3**

**Supplemental Figure 1 only for review, related to Table 1.**

**Supplemental data 1 for Fig. 2B**

**Supplemental data 2 for Fig. 2C**

**Supplemental data 3 for Fig. 3B**

**Supplemental data 4 for Fig. 3I**

**Supplemental data 5 for Fig. 4A**

**ONLINE EXPERIMENTAL PROCEDURES**

***DLBCL tissue samples, cell lines and reagents***

We examined the medical history of all DLBCL patients from 2008 to 2015 at Fudan University Shanghai Cancer Center and found a total of 12 patients who simultaneously had both paraffin-embedded tissue samples from the initial visit and from relapse. DLBCL cases were subgrouped into GCB (6 cases) or ABC (6 cases) molecular subtypes based on the Hans immunohistochemistry algorithm. The relapsed patients received R-CHOP therapy for at least 6 cycles. The first biopsy was performed for diagnosis at the initial visit, and the second biopsy was performed to detect the relapsed status or to confirm the primary diagnosis. The interval of the two biopsies differed from one another among the patients. These DLBCL samples were subsequently collected for IHC staining. The Ethics Committee at Fudan University Shanghai Cancer Center approved the study, and all patients signed the informed consent form.

The human DLBCL cell lines GCB subtype OCI-LY8 and ABC subtype NU-DUL-1 were a kind gift from Professor Xiaoyan Zhou, Department of Pathology, Fudan University Shanghai Cancer Center (Shanghai, China). LY8 cells were maintained in Iscove's modified Dulbecco's medium supplemented with 10% fetal bovine serum and 1% penicillin/streptomycin. NU-DUL-1 cells were maintained in RPMI 1640 medium supplemented with 10% fetal bovine serum and 1% penicillin/streptomycin. The results of STR analysis for LY8 (OCI-Ly8) and NU-DUL-1 cell lines are showed in the following table:

Results of cell line fingerprinting using STR.

| **Source** | **Sample**  **Name** | **D5S818** | **D13S317** | **D7S820** | **D16S539** | **VWA** | **TH01** | **AM** | **TPOX** | **CSF1PO** | **Comment** |
| --- | --- | --- | --- | --- | --- | --- | --- | --- | --- | --- | --- |
| Present study | OCI-LY8 | 10,13 | 11,11 | 11,12 | 9,11 | 17,19 | 6,9.3 | X,X | 9,11 | 11,12 | MATCH |
| PMID: 23127183 | OCI-LY8 | 10,13 | 11,11 | 11,12 | 9,11 | 17,19 | 6,9.3 | X,X | 9,11 | 11,12 |  |
| Present study | NU-DUL-1 | 12,14 | 8,13 | 11,13 | 11,13 | 16,17 | 9,9.3 | X,X | 8,8 | 11,12 | MATCH |
| ATCC | NU-DUL-1 | 12,14 | 8,13 | 11,13 | 11,13 | 16,17 | 9,9.3 | X,X | 8,8 | 11,12 |  |

As a complement source, normal human serum (NHS) was pooled from 10 healthy persons, aliquoted and stored at −80°C until use. Heat-inactivated human serum (IHS) as a negative control was prepared in a 65°C water bath for 30 minutes before use.

The anti-SOX2-T118p and anti-SOX2-K119me polyclonal antibodies were generated previously (*1*). Information on the commercial antibodies used in this study is provided in Table S1.

The PI3K inhibitor IPI-145 (duvelisib), BAY80-6949 (copanlisib), CDK6 inhibitor abemaciclib, FGFR1/2 inhibitor AZD4547, FAK inhibitor PF-573228, Syk inhibitor R788, Src inhibitor saracatinib, Verapamil hydrochloride, Hoechst 33342 and prednisolone were purchased from MedChem Express (Monmouth Junction, NJ).

***Generation and characterization of RCHO-resistant DLBCL cells***

We generated LY8 and NU-DUL-1 cells that were resistant to rituximab-mediated complement-dependent cytotoxicity (CDC), as previously described (*2*). In brief, original LY8 (LY8-ORI) and NU-DUL-1 (NU-DUL-1-ORI) cells were treated with escalating rituximab (Roche, Basel, Switzerland) concentrations from 4 μg/mL to 32 μg/mL in the presence of 20% NHS. The resistant cells were denoted LY8-R and NU-DUL-1-R. LY8-R and NU-DUL-1-R cells were treated with 32 µg/mL rituximab and 20% NHS every 21 days to maintain resistance. The cytolysis induced by CDC was assessed by detecting propidium iodide (PI) staining-positive cells with a fluorescence-activated cell sorting (FACS) assay.

Chemo-resistant LY8 and NU-DUL-1 cells were generated as previously described(*3*). Briefly, LY8-ORI and NU-DUL-1-ORI cells were treated with doxorubicin (Selleck Chemicals, Houston, TX) and vincristine (Selleck Chemicals, Houston, TX) at the clinical ratio of 50:1.4 by escalating the concentration. The maximum resistant dosage for LY8 cells was 125 ng/mL doxorubicin and 3.5 ng/mL vincristine, whereas it was 25 ng/mL doxorubicin and 0.7 ng/mL vincristine for NU-DUL-1 cells. These cells were then treated with 2 µg/mL 4-hydroperoxycyclophosphamide (4-HC) (Santa Cruz Biotechnology, Santa Cruz, CA) every 21 days for 3 cycles. The obtained CHO-resistant cells were termed LY8-CHO and NU-DUL-1-CHO, respectively. The CHO-resistant cells were treated with doxorubicin, vincristine and 4-HC every 21 days to maintain CHO resistance.

Using a similar approach, we treated the LY8-R and NU-DUL-1-R cells to generate LY8-RCHO-resistant cells and NU-DUL-1-RCHO-resistant cells. To maintain RCHO resistance, the LY8-RCHO and NU-DUL-1-RCHO cells were treated with doxorubicin, vincristine and 4-HC every 21 days following treatment with 32 µg/mL rituximab and 20% NHS. CHO and RCHO resistance were determined by serial CCK-8 assays after doxorubicin, vincristine and 4-HC treatment for 48 hours.

***Aldefluor Assay***

ALDH1 is a selectable marker for multiple kinds of normal and cancer stem cells, including hematopoietic stem cells (*4, 5*). Thus, we evaluated tumor propagating cell numbers in hematopoietic malignancies by detecting ALDH1-positive cells. We employed an ALDEFLUOR™ kit (StemCell Technologies, Vancouver, BC, CA) to detect populations with high ALDH1 activity as previously described. Briefly, 1×10^6^ cells were suspended in 1 mL of assay buffer containing 1 µM ALDH1 substrate BAAA. The suspended cells were then equally divided into two aliquots, one as the negative control and the other as the test sample. To form the negative control, 50 mM DEAB, a specific ALDH inhibitor, was added to one aliquot. The cells were then incubated for 30 minutes at 37°C before flow cytometry analysis.

For concurrently detecting CD133 population and ALDH1 activity, the RCHO-resistant cells were incubated with Human TruStain FcX™ (Fc Receptor Blocking Solution) for 30 minutes at room temperature, then stained with anti-CD133 antibody for 30 minutes before the procedures of Aldefluor Assay.

***Sphere Formation Assay***

We conducted sphere formation assays as previously described (*6*). In brief, cells were suspended in serum-free medium (DMEM/F12, 3:1 mixture) containing 0.4% BSA and 0.2× B27 lacking vitamin A (Life Technologies, Gaithersburg, MD) and supplemented with recombinant EGF (PeproTech, Rocky Hill, NJ) at 10 ng/mL, recombinant basic fibroblast growth factor (PeproTech, Rocky Hill, NJ) at 10 ng/mL and insulin (Sigma-Aldrich, St. Louis, MO) at 5 μg/mL. The cells were then seeded in ultra-low attachment 24-well plates at a density of 1×10^4^ cells/mL. The medium was replaced every 7 days, and the spheres were counted and harvested at 14 days. The sphere cells were subcultured with the above medium at clonal density. Images were captured on the 14th day of cultivation after 3 passages.

***CytoTox-Glo™ Cytotoxicity Assay***

Cells were plated in 96-well plates at a density of 1×10^4^ cells/100 µL/well. The cells were pretreated with IPI-145, abemaciclib, AZD4547, PF-573228, R788, or saracatinib at escalating concentrations in the presence or absence of CHO for 48 h before performing the CytoTox-Glo™ cytotoxicity assays. We used a CytoTox-Glo™ cytotoxicity assay kit (Promega, Madison, WI) to perform these assays according to the technical bulletin. Briefly, 50 µL of CytoTox-Glo™ cytotoxicity assay reagent was added to all wells, mixed by orbital shaking and then incubated for 15 minutes at room temperature. Experimental dead cell luminescence was measured with a Synergy HT microplate reader (BioTek, Biotek Winooski, MN), in which 50 µL of lysis reagent was added to all wells, mixed and incubated at room temperature for 15 minutes, followed by measurement of total luminescence. Cytotoxicity was calculated according to the following formula: Cytotoxicity (%) = Experimental dead cell luminescence/Total luminescence × 100%.

***Immunohistochemistry***

Tumor tissues derived from patients or animal models were fixed with 4% formalin, embedded in paraffin and sectioned. Paraffin sections were incubated with 3% hydrogen peroxide to block endogenous peroxidase for 15 minutes at 37°C and rinsed with 0.01 M PBS, followed by high-pressure antigen retrieval in EDTA buffer. The sections were then incubated with rabbit anti-SOX2 monoclonal antibody (1:200; Cell Signaling Technology, Danvers, MA) at 4°C overnight. After rinsing 3 times in PBS, the sections were incubated with peroxidase-conjugated AffiniPure goat anti-rabbit IgG H&L (1:200; Proteintech, Chicago, IL) at room temperature for 1 hour. Then, immunoreactivity was measured using a GTVision III immunohistochemical detection kit (GK500705; Gene Tech, Shanghai, China) according to the manufacturer’s instructions. The dilution ratio of primary antibodies for immunohistochemical staining of tissue microarrays is shown in Table S1. The immunostaining scores for SOX2, p-AKT, CDK6, FGFR1 and FGFR2 were assessed under a microscope by three independent individuals according to the following formula: score = 3 (strong positive) × percentage + 2 (moderate positive) × percentage + 1 (weak positive) × percentage + 0 (negative) × percentage (*7*).

***Immunoblotting Assay***

We performed immunoblotting assays according to the standard protocol, and the related antibodies are shown in Table S1.

***Quantitative Real-time PCR (qRT-PCR)***

We performed qRT-PCR as previously described (*6*). Briefly, total RNA from cells was extracted with TRIzol reagent (Invitrogen, Grand Island, NY) and then transcribed into cDNA using a Reverse Transcription System (Promega, Madison, WI). The input cDNA was standardized and amplified for 40 cycles with SYBR Green Master Mix (Invitrogen, Grand Island, NY) and gene-specific primers on a Roche LightCycler 480 system (Roche, Basel, Switzerland). We used the *ACTB* gene encoding β-actin as the endogenous control, and the samples were analyzed in triplicate. The primers for qRT-PCR are listed in Table S2.

***FACS Analysis***

After washing with PBS, cells were incubated with fluorescein-conjugated antibodies for 30 minutes and then rinsed and resuspended in PBS. Fluorochrome-conjugated isotype-specific IgGs served as controls. Flow cytometric analysis was performed on a Cytomics FC500 MPL instrument (Beckman Coulter, Brea, CA) and analyzed with FlowJo software (Ashland, OR). We performed cell sorting with a MoFlo XDP instrument (Beckman Coulter, Brea, CA) according to the relative fluorescence.

For concurrently detecting SOX2 and FGFR1/2, the RCHO-resistant cells were stained with anti-FGFR1 or anti-FGFR2 antibody for 30 minutes before using the True-Nuclear™ Transcription Factor Buffer Set (BioLegend, San Diego, CA) for fixation and permeabilization. Then the cells were stained with anti-SOX2 antibody for 30 minutes and the related secondary antibodies conjugated with different fluorescence.

For concurrently detecting SOX2 and p-AKT1 (S473), CDK6, the RCHO-resistant cells were fixed and permeabilized using the True-Nuclear™ Transcription Factor Buffer Set (BioLegend, San Diego, CA). Then the cells were stained with anti-SOX2 and anti-p-AKT1 (S473) or anti-CDK6 antibodies for 30 minutes, subsequently with related secondary antibodies conjugated with different fluorescence.

***Side Population Assay***

The side population (SP) of the original and resistant DLBCL cells were identified using the DNA-binding dye Hoechst 33342 (MedChem Express, Monmouth Junction, NJ) and a MoFlo XDP instrument (Beckman Coulter, Brea, CA) based on the protocol described by Goodell et al (*8*). Briefly, a total of 5 x 10^7^ cells were harvested by centrifugation, and re-suspended with pre-warmed DMEM medium containing 2% FBS. Then the cells were stained with Hoechst 33342 in a final concentration of 5 µg/mL in the 37°C water bath in the dark, either alone or in the present of 100 µM Verapamil hydrochloride (MedChem Express, Monmouth Junction, NJ). After two-hour incubation, the cells were centrifugated and re-suspended with pre-cold HBSS buffer containing 2% FBS and 10mM Hepes. Finally, 1 µg/mL propidium iodine (PI) (Sigma-Aldrich, St. Louis, MO) was added for the discrimination of dead cells. The cells were maintained on the ice until analysis.

***EdU Cell Proliferation Assay***

For EdU assays on SOX2-positive cells, we used the True-Nuclear™ Transcription Factor Buffer Set (BioLegend, San Diego, CA) for fixation and permeabilization. The RCHO-resistant cells were subsequently stained with anti-SOX2 antibody conjugated with APC fluorescence for 30 minutes. Then EdU proliferation assay were conducted according to the manufacturer’s instructions of the Cell-Light EdU Apollo488 In Vitro Flow Cytometry Kit (Guangzhou RiboBio Co., Ltd., Guangzhou, China). Next, we assessed EdU staining-positive cells on a Cytomics FC500 MPL instrument (Beckman Coulter, Brea, CA).

***Plasmid Construction and Lentiviral Transduction***

The coding DNA sequence (CDS) of human *SOX2* was obtained by PCR amplification from cDNA pools of A549 cells as previously described (*6*). This sequence was cloned into the pCDH cDNA cloning and expression lentivector (cat# CD511B-1, System Biosciences, Palo Alto, CA 94303) via EcoRI and BamHI endonuclease sites for stable *SOX2* overexpression in OCI-LY8 and NU-DUL-1 cells. The CDS of the *Firefly Luciferase* gene was obtained by PCR amplification from the pGL3-Basic plasmid and inserted into the pCDH cDNA cloning and expression lentivector via EcoRI and BamHI endonuclease sites. The CDS of human *AKT1* with a myristoylation signal (termed *myr-AKT1*) was obtained by adding the myristoylation signal coding sequence to the forward primer during the PCR amplification from cDNA pools of OCI-LY8 cells. Then this sequence was inserted into the pCDH cDNA cloning and expression lentivector via EcoRI and BamHI endonuclease sites. The CDS of human *OCT4* was obtained by PCR amplification from cDNA pools of OCI-LY8 cells, and inserted into the pCDH cDNA cloning and expression lentivector via EcoRI and BamHI endonuclease sites. The pLKO.3G cloning vector (plasmid #14748, Addgene, Cambridge, MA) was employed to construct scramble (SCR), *SOX2*, *ITGA1*, *ITGB5*, *CD79A*, and *CCR7* shRNA plasmids. 293FT cells were co-transfected with the pCDH or pLKO.3G plasmid and pMD.2G and psPAX2 plasmids to generate *SOX2*, *OCT4*, *myr-AKT1*, *firefly luciferase* overexpression or *SOX2*, *ITGA1*, *ITGB5*, *CD79A*, or *CCR7* knockdown lentivirus, respectively. The lentivirus was subsequently added to OCI-LY8, NU-DUL-1, LY8-RCHO or NU-DUL-1-RCHO culture medium for 48 hours of incubation. All the cells transduced with lentivirus in this study were sorted by GFP with a MoFlo XDP instrument (Beckman Coulter, Brea, CA). Information regarding the shRNA oligonucleotide sequences is shown in Table S2.

***Xenograft Model***

To compare the tumor-initiating capacity of RCHO-resistant and original cells, six-week-old male NOD-SCID mice were purchased from Slac Laboratory Animal Center (Shanghai, China). Then, two pairs of cells, LY8-ORI and LY8-RCHO cells, and NU-DUL-1-ORI and NU-DUL-1-RCHO cells, at the dose of 5x10^2^, 5x10^3^, 5x10^4^, 5x10^5^, or 5x10^6^ cells were re-suspended in a PBS/Matrigel (Invitrogen, Grand Island, NY) mixture (1:1 volume). Further, each dose of RCHO-resistant or original cells were injected subcutaneously into bilateral backside, respectively, in same mouse, 5 mice in for each dose. The mice were monitored for tumor volume and incidence. The tumor volume was determined following a standard formula: Length×Width^2^/2. The tumor-free mice were followed up until 80 days after implantation.

To test the drug efficacy on suppressing tumor growth, we transduced the LY8-RCHO and NU-DUL-1-RCHO cells with lentivirus overexpressing the *firefly luciferase* gene to generate stable cells expressing firefly luciferase, termed LY8-RCHO-Luc and NU-DUL-1-RCHO-Luc cells, respectively. Eight-week-old female SCID mice were purchased from Slac Laboratory Animal Center (Shanghai, China). LY8-RCHO-Luc and NU-DUL-1-RCHO-Luc cells were resuspended in PBS and then intraperitoneally injected at 1.5×10^7^ cells per mouse. The mice were randomly divided into 6 groups (7 mice per group) after intraperitoneal injection with LY8-RCHO-Luc cells and administered saline, R-CHOP, duvelisib, abemaciclib, R-CHOP+duvelisib, or R-CHOP+abemaciclib. The mice bearing NU-DUL-1-RCHO-Luc cells were also randomly divided into 6 groups (7 mice per group), which were administered saline, R-CHOP, duvelisib, AZD4547, R-CHOP+duvelisib, or R-CHOP+AZD4547. The methods of drug delivery based on the clinical usage for one cycle are indicated in Table S3; the R-CHOP regimen was employed in clinical use for DLBCL therapy (*9*), and duvelisib, abemaciclib, and AZD4547 were tested in clinical trials (the related NCT numbers are NCT02576275, NCT01739309, and NCT01739309, respectively). For the first time of drug delivery, duvelisib, abemaciclib and AZD4547 were administrated 8 hours before R-CHOP delivery. Tumor growth was monitored by bioluminescence at 50 and 90 days after implantation. For in vivo luminescence imaging, D-luciferin (Promega, Madison, WI) was injected intraperitoneally into the mice (150 mg/kg). After 10 minutes, the mice were anesthetized by intraperitoneal injection with pentobarbital (50 mg/kg), and then bioluminescence was examined using an In Vivo MS FX PRO system (Bruker, Billerica, MA). Luminescence images were captured with 30 seconds of exposure time, and the signal intensities of the tumors were measured using Bruker MI software. The survival time of each mouse was recorded. The surviving mice were euthanized by CO_2_ and dissected at 120 days after xenografting, and no intraperitoneal tumors were found. Tumor tissues were collected from the moribund mice euthanized by CO_2_ and fixed with 4% formalin.

All the animal experiments were conducted in strict accordance with experimental protocols approved by the Animal Ethics Committee at Shanghai Medical School, Fudan University.

***RNA Sequencing and Bioinformatic Analysis***

Total RNA was extracted from LY8-ORI, LY8-R, LY8-CHO, LY8-RCHO, NU-DUL-1, NU-DUL-1-R, NU-DUL-1-CHO, and NU-DUL-1-RCHO cells with TRIzol reagent (Invitrogen, Grand Island, NY). The total RNA from each group from 3 different passages was pooled separately. RNA sequencing (RNA-seq) and bioinformatics analysis were conducted by Shanghai Novelbio Ltd. according to their established procedures (*10*). We applied the DEseq algorithm to filter the differentially expressed genes after significance and false discovery rate (FDR) analyses under the following criteria: (1) fold change >1.5 or <0.667; (2) FDR<0.05. Pathway analysis was used to identify significant pathways of the differential genes according to the KEGG database. Series-cluster analysis was performed to identify global trends and model profiles of expression according to the signal density of the ORI-R-RCHO and ORI-CHO-RCHO sequences. Fisher’s exact test and the multiple comparison test were used to select significant pathways, and the threshold of significance was defined by *P*-value and FDR (*11, 12*). We screened significantly enriched pathways with escalating trends in ORI-R-RCHO and ORI-CHO-RCHO sequences and then analyzed the interaction frequency of these pathways between these groups using Cytoscape software (*13*).

We performed gene set enrichment analysis (GSEA) to identify functions of differentially expressed genes based on RNA-seq using GSEA software from the Broad Institute (Massachusetts Institute of Technology) (*14, 15*). The pre-ranked version of the software was used to identify significantly enriched pathways, and pathways enriched with FDR<0.25 were considered significant. The PI3K-AKT signaling pathway gene set used in this study consisted of 341 genes from the PI3K-AKT signaling pathway SuperPath in the PathCards pathway unification database (Version 4.6.0.37, Weizmann Institute of Science). The GSEA term KEGG_CHEMOKINE_SIGNALING_PATHWAY was also used to identify the chemokine signaling pathway.

The RNA-seq data reported in this study are accessible in the NCBI GEO database through GEO series accession numbers GSE112989 and GSE113001.

***Statistical Methods***

The data are presented as mean ± SD unless otherwise specified. Significant differences between two groups were determined using two-tailed Student’s t-test for unpaired data, and *P*<0.05 was considered statistically significant. For IHC staining scores of tissue microarrays, significance was determined by two-tailed paired t-test, and *P*<0.05 was considered statistically significant. For the total photon flux of animal models, significance was determined by a one-tailed Mann-Whitney test, and *P*<0.05 was considered statistically significant. We applied a Mantel-Cox test to compare the survival rates between the two groups of xenograft models, and *P*<0.05 was considered statistically significant. All statistical analyses were performed by GraphPad Prism 7 (La Jolla, CA).

**References**

1.	L. Fang *et al.*, A methylation-phosphorylation switch determines Sox2 stability and function in ESC maintenance or differentiation. *Molecular cell* **55**, 537-551 (2014).

2.	W. Hu *et al.*, Human CD59 inhibitor sensitizes rituximab-resistant lymphoma cells to complement-mediated cytolysis. *Cancer Res* **71**, 2298-2307 (2011).

3.	S. A. Maxwell, E. M. Cherry, K. J. Bayless, Akt, 14-3-3zeta, and vimentin mediate a drug-resistant invasive phenotype in diffuse large B-cell lymphoma. *Leukemia & lymphoma* **52**, 849-864 (2011).

4.	C. Ginestier *et al.*, ALDH1 is a marker of normal and malignant human mammary stem cells and a predictor of poor clinical outcome. *Cell stem cell* **1**, 555-567 (2007).

5.	J. P. Chute *et al.*, Inhibition of aldehyde dehydrogenase and retinoid signaling induces the expansion of human hematopoietic stem cells. *Proc Natl Acad Sci U S A* **103**, 11707-11712 (2006).

6.	J. Chen *et al.*, CD59 Regulation by SOX2 Is Required for Epithelial Cancer Stem Cells to Evade Complement Surveillance. *Stem Cell Reports* **8**, 140-151 (2017).

7.	X. Zhang *et al.*, A novel FOXM1 isoform, FOXM1D, promotes epithelial-mesenchymal transition and metastasis through ROCKs activation in colorectal cancer. *Oncogene* **36**, 807-819 (2017).

8.	M. A. Goodell, K. Brose, G. Paradis, A. S. Conner, R. C. Mulligan, Isolation and functional properties of murine hematopoietic stem cells that are replicating in vivo. *J Exp Med* **183**, 1797-1806 (1996).

9.	D. Cunningham *et al.*, Rituximab plus cyclophosphamide, doxorubicin, vincristine, and prednisolone in patients with newly diagnosed diffuse large B-cell non-Hodgkin lymphoma: a phase 3 comparison of dose intensification with 14-day versus 21-day cycles. *Lancet* **381**, 1817-1826 (2013).

10.	Z. Liu *et al.*, Autism-like behaviours and germline transmission in transgenic monkeys overexpressing MeCP2. *Nature* **530**, 98-102 (2016).

11.	M. F. Ramoni, P. Sebastiani, I. S. Kohane, Cluster analysis of gene expression dynamics. *Proc Natl Acad Sci U S A* **99**, 9121-9126 (2002).

12.	L. D. Miller *et al.*, Optimal gene expression analysis by microarrays. *Cancer Cell* **2**, 353-361 (2002).

13.	P. Shannon *et al.*, Cytoscape: a software environment for integrated models of biomolecular interaction networks. *Genome Res* **13**, 2498-2504 (2003).

14.	V. K. Mootha *et al.*, PGC-1alpha-responsive genes involved in oxidative phosphorylation are coordinately downregulated in human diabetes. *Nat Genet* **34**, 267-273 (2003).

15.	A. Subramanian *et al.*, Gene set enrichment analysis: a knowledge-based approach for interpreting genome-wide expression profiles. *Proc Natl Acad Sci U S A* **102**, 15545-15550 (2005).

**Supplemental Figures and Legends**

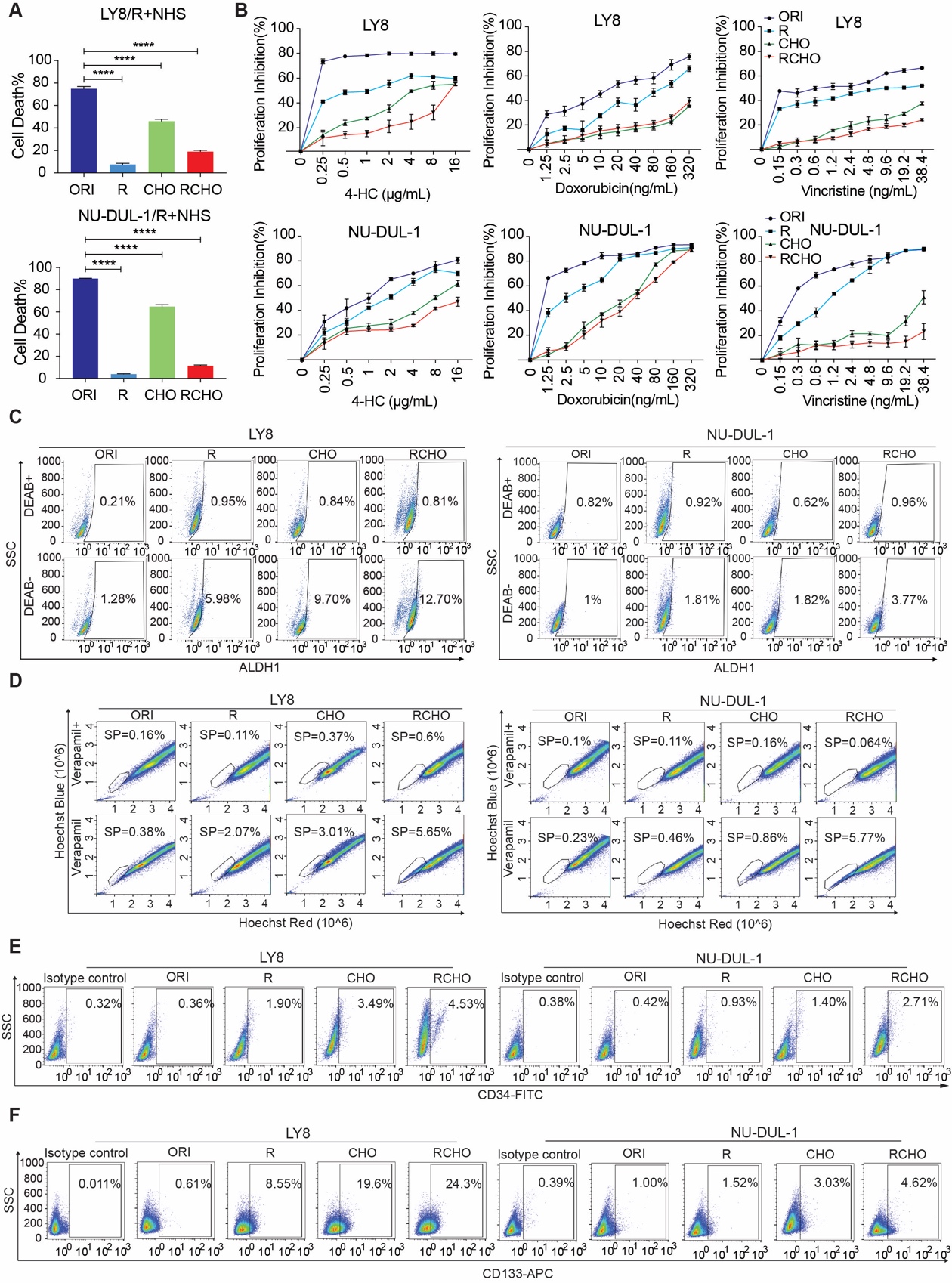

**Fig. S1. Resistance validation and stemness characterization of the resistant DLBCL cells/tissues.** (**A and B**) Resistance to R-mediated CDC (A) or chemotherapy (B) was confirmed by a CDC assay (A) or CCK-8 assay (B), respectively, in which the cells were treated with 3.2 µg/mL rituximab and 20% NHS for 2 hours (A) or by escalating dosages of 4-HC, doxorubicin, or vincristine for 48 hours (B). (**C**) Representative images of the Aldefluor assay. The percentage of ALDH1-positive cells in resistant DLBCL cells was gradually enhanced in order of R-, CHO- and RCHO-resistant cells compared with the original cells, indicating increased stemness of the resistant DLBCL cells. The related quantitative results are shown in Fig. 1C by subtracting the DEAB controls. (**D**) Representative images of the side population assay. The percentage of side population fraction in resistant DLBCL cells was gradually increased in order of R-, CHO- and RCHO-resistant cells compared with the original cells. The related quantitative results are shown in Fig. 1D by subtracting the Verapamil control. (**E and F**) Representative images of CD34 and CD133 detection by flow cytometry. Fluorochrome-conjugated isotype-specific IgGs were used to set the gate for CD34- or CD133-positive cells. The percentage of CD34- and CD133-positive cells in resistant DLBCL cells was gradually increased in order of R-, CHO- and RCHO-resistant cells compared with the original cells. The related quantitative results are shown in Fig. 1E and F. The data are presented as mean ± SD; n=3. *****P*<0.0001.

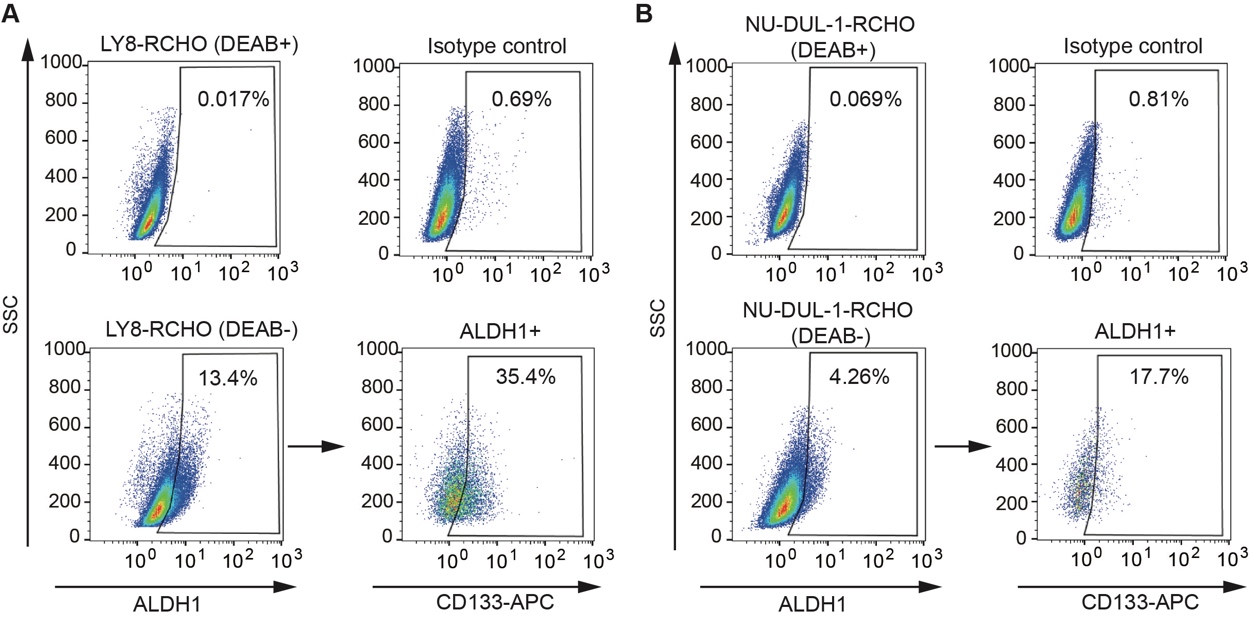

**Fig. S2. Overlapped population of ALDH1^+^ and CD133^+^ cells.** (**A**-**B**) Representative images for detecting the overlapped population of ALDH1^+^ and CD133^+^ cells in RCHO-resistant DLBCL cells by flow cytometry. The DEAB controls were used for gating the ALDH1^+^ cells, and the Fluorochrome-conjugated isotype-specific IgGs were used for gating the CD133^+^ cells.

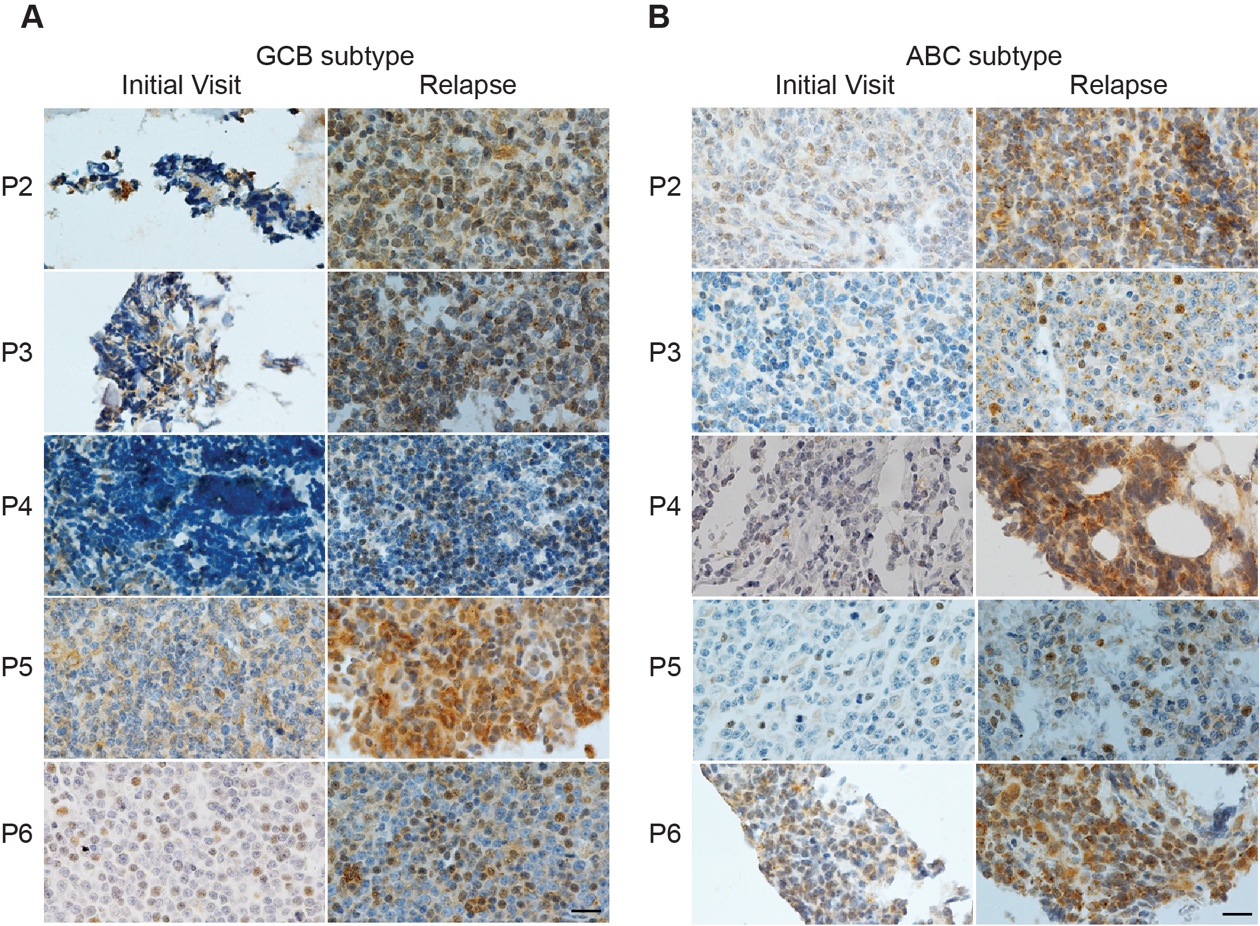

**Fig. S3. SOX2 levels in clinical samples.** (**A**-**B**) SOX2 expression detected by IHC staining was markedly increased in relapsed 6 GCB (A) and 6 ABC (B) subtype DLBCL tissues compared with the paired patient tissues from the initial visit. Image from patient 1 and quantitative results for the GCB and ABC subtypes are shown in Fig. 1I. P: patient. Scale bar: 20 μm.

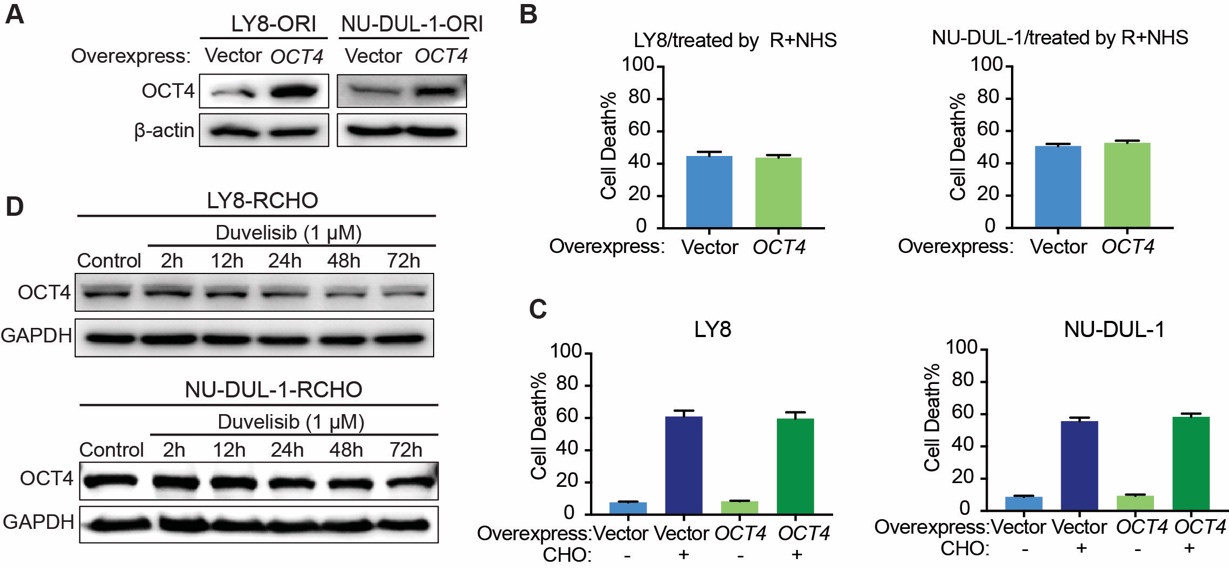

**Fig. S4. The effect of ectopic OCT4 expression in original cells on resistance to R and CHO treatment.** (**A**) Validation of ectopic OCT4 expression in original DLBCL cells by immunoblotting. (**B**) CDC assays: ectopic OCT4 expression failed to alter the susceptibility of original cells to R-mediated CDC. (**C**) CytoTox-Glo cytotoxicity assays: ectopic OCT4 expression in original cells failed to elevate the resistance to CHO. (**D**) OCT4 expression reduced after duvelisib treatment for 24 hours in RCHO-resistant LY8 (up) and NU-DUL-1 (bottom) cells.

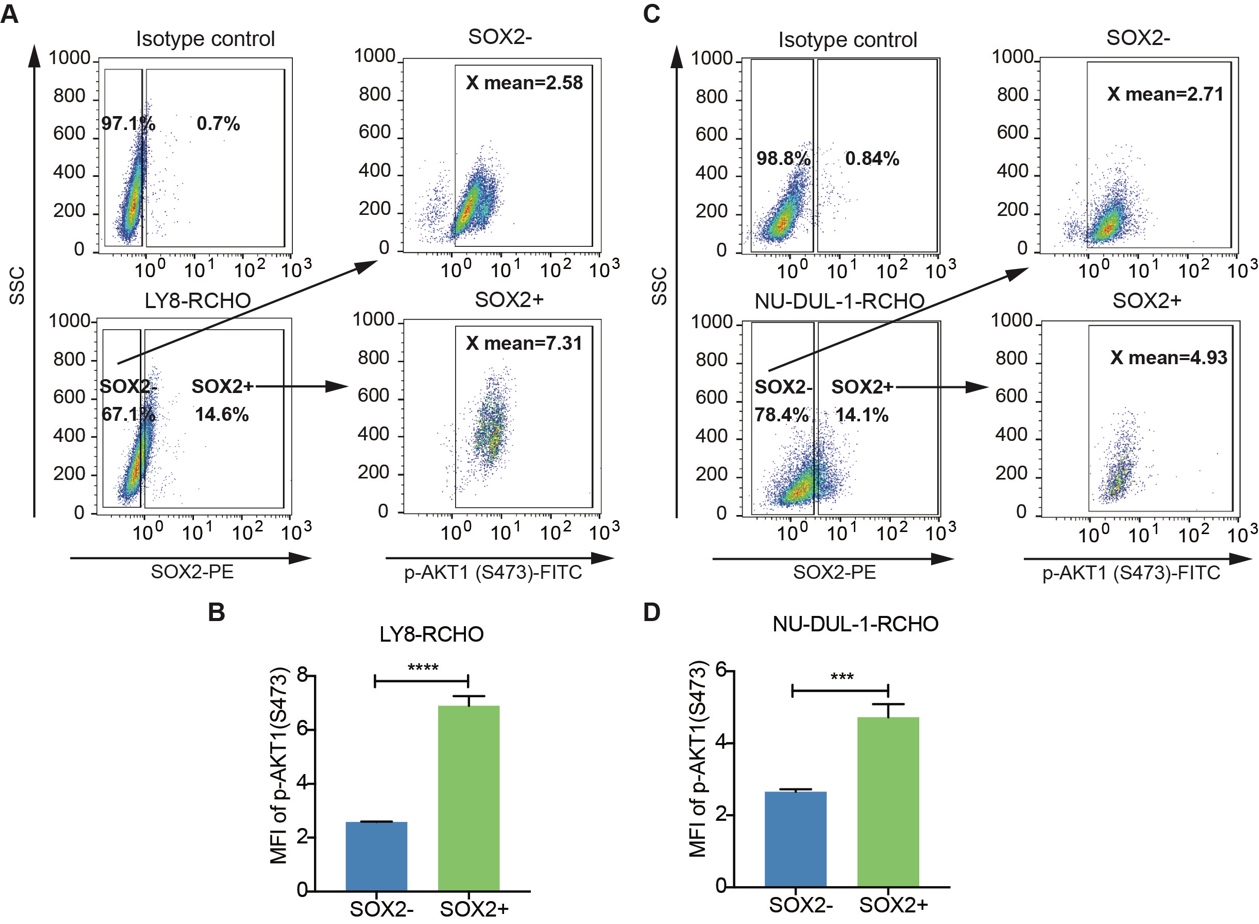

**Fig. S5. SOX2^+^ subpopulation displayed higher AKT activation.** (**A**-**D**) Representative images (A and C) for detecting p-AKT1 (S473) levels in SOX2^-^ and SOX2^+^ subpopulations of RCHO-resistant LY8 (A) and NU-DUL-1 (C) cells. The Fluorochrome-conjugated isotype-specific IgGs were used for gating the SOX2^+^ cells and p-AKT1 (S473) positive cells. The quantitative results showed that SOX2^+^ subpopulation exhibited higher p-AKT1 (S473) level than SOX2^-^ subpopulation in RCHO-resistant LY8 (B) and NU-DUL-1 (D) cells. The data are presented as mean ± SD; n=3. ****P*<0.001, *****P*<0.0001.

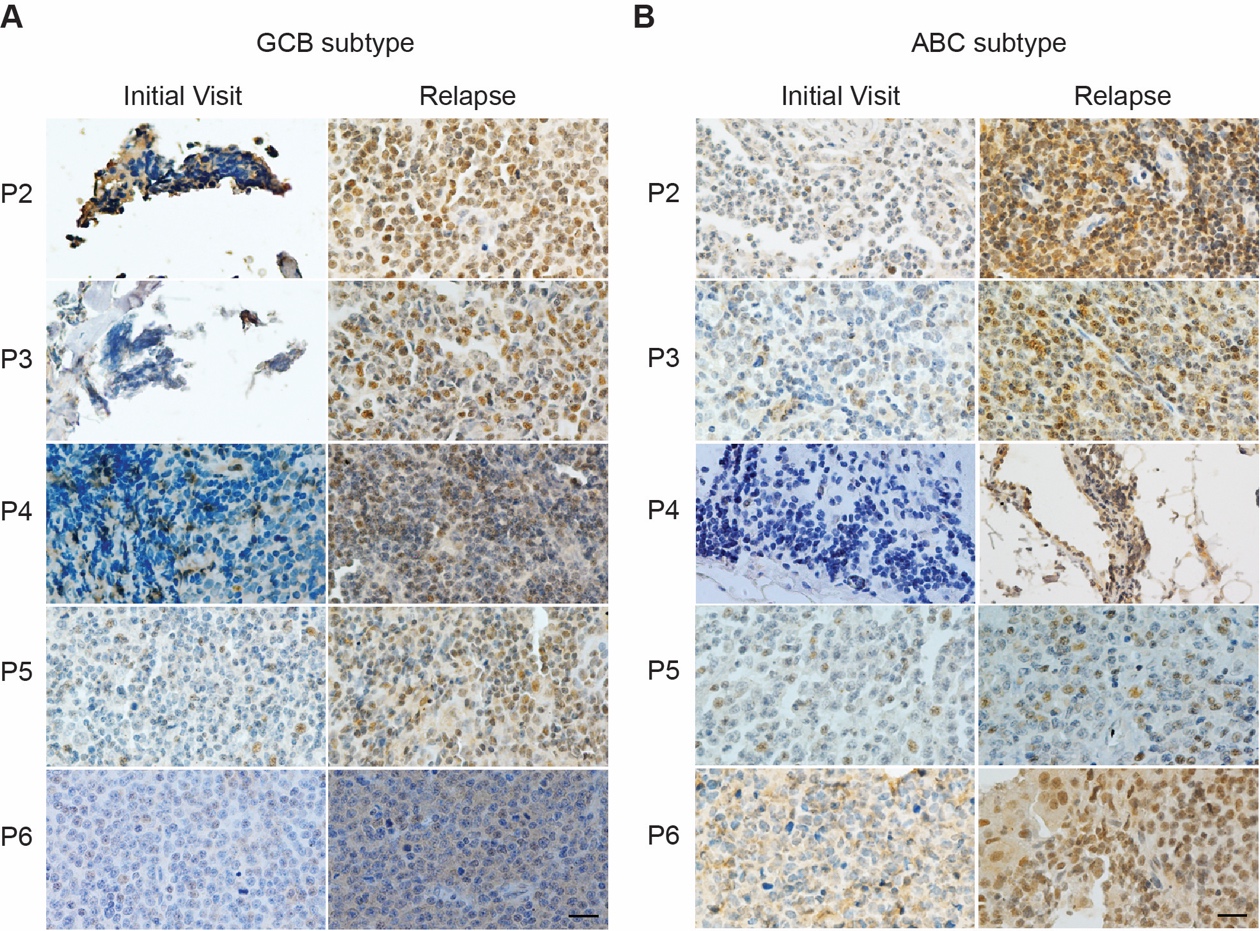

**Fig. S6. AKT activation in relapsed DLBCL tissues.** (**A**-**B**) AKT activation determined by IHC staining for p-AKT (S473) was significantly enhanced in relapsed 6 GCB (A) and 6 ABC (B) subtype DLBCL tissues compared with the paired patient tissues from the initial visit. Image from patient 1 and quantitative results for the GCB and ABC subtypes are shown in Fig. 2E. P: patient. Scale bar: 20 μm.

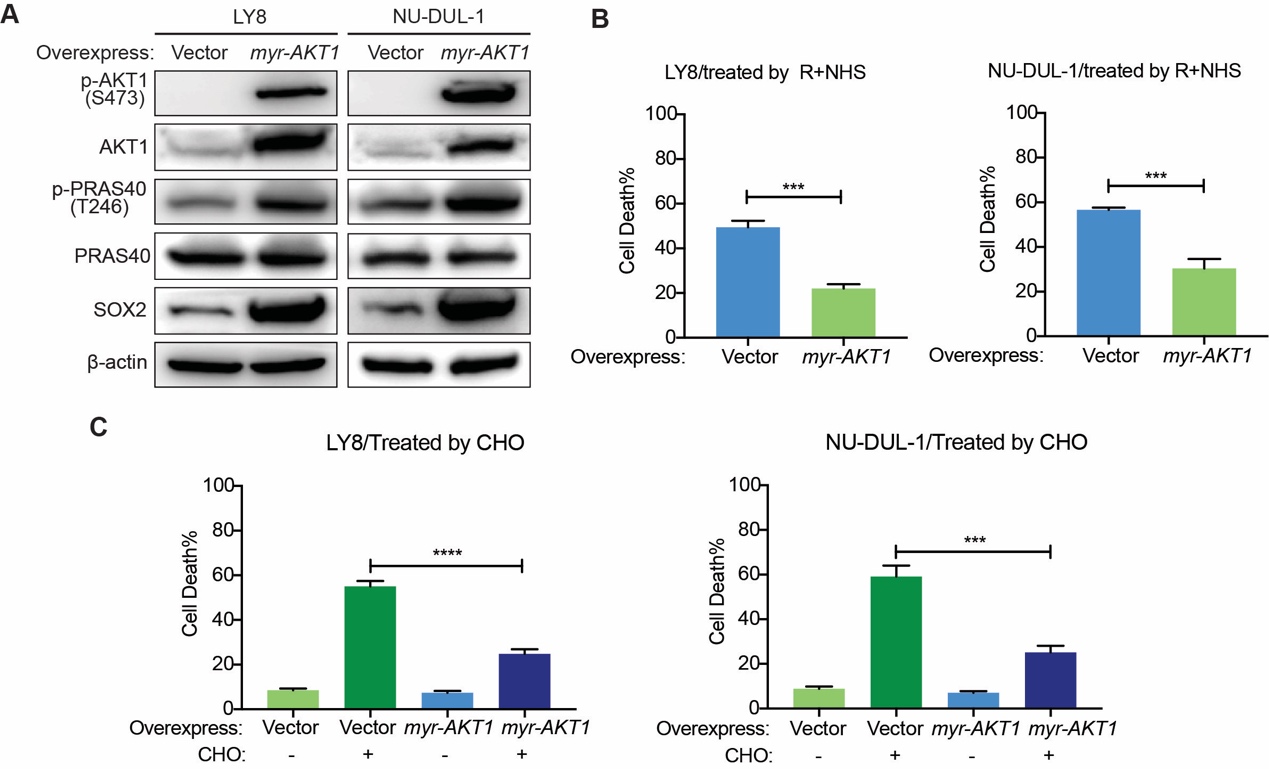

**Fig. S7. Ectopic myr-AKT1 expression** **in original DLBCL cells increased the resistance to R and CHO treatment by up-regulating SOX2.** (**A**) Ectopic myr-AKT1 expression resulted in constitutive AKT activation and elevated the levels of p-PRAS40 and SOX2. (**B**) CDC assays: ectopic myr-AKT1 expression reduced the susceptibility to R-mediated CDC. (**C**) CytoTox-Glo cytotoxicity assays: ectopic myr-AKT1 expression elevated the resistance to CHO treatment. The data are presented as mean ± SD; n=3. ****P*<0.001, and *****P*<0.0001.

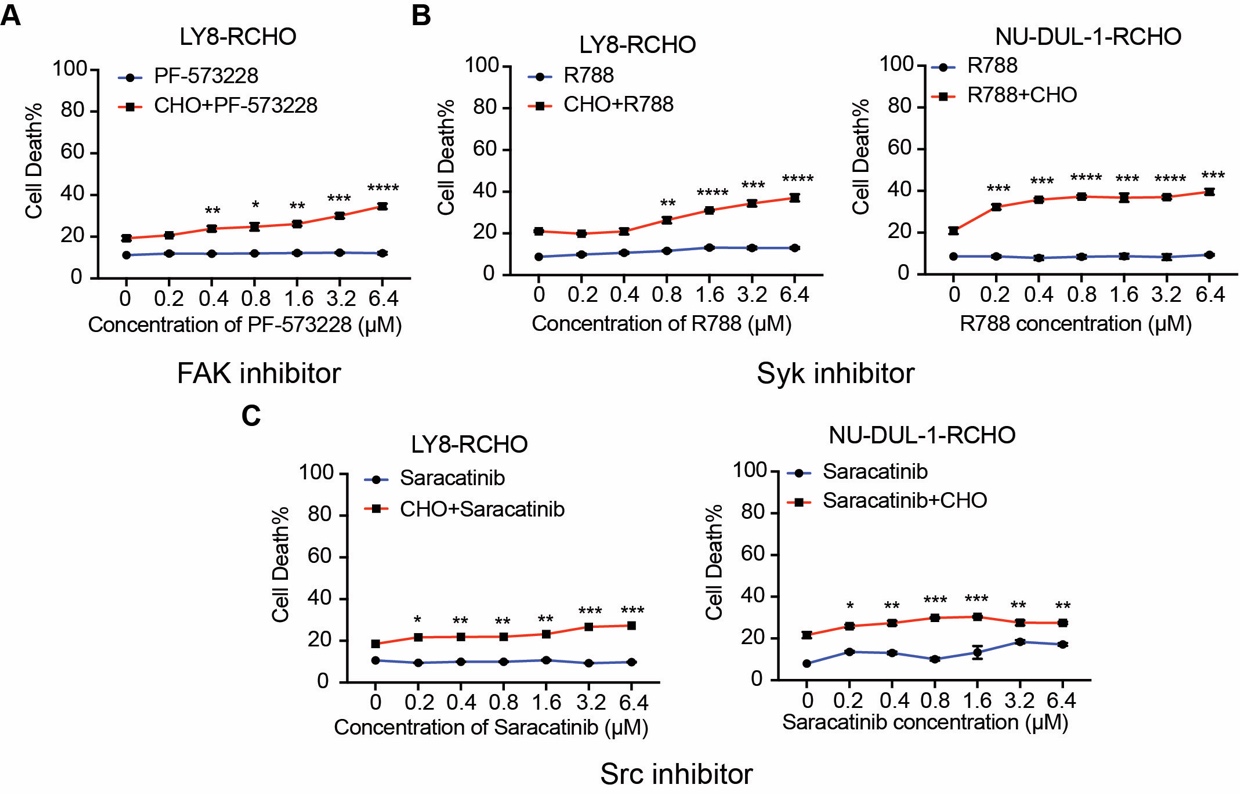

**Fig. S8. Inhibition of FAK, Syk or Src reversed resistance to CHO treatment in RCHO-resistant DLBCL cells.** (**A**-**C**) Addition of FAK (A), Syk (B), or Src (C) inhibitors to CHO reversed resistance to CHO in the indicated RCHO-resistant DLBCL cells to a certain degree. Inhibitors alone exhibited a negligible cytotoxic effect on the induction of cell death. A CytoTox-Glo cytotoxicity assay was employed to measure the cytotoxic effect. The cells were treated with PF-573228, R788, or saracatinib in the presence or absence of CHO for 48 hours before assays. The data are presented as mean ± SD; n=3. **P*<0.05, ***P*<0.01, ****P*<0.001 and *****P*<0.0001 *vs* CHO alone treatment.

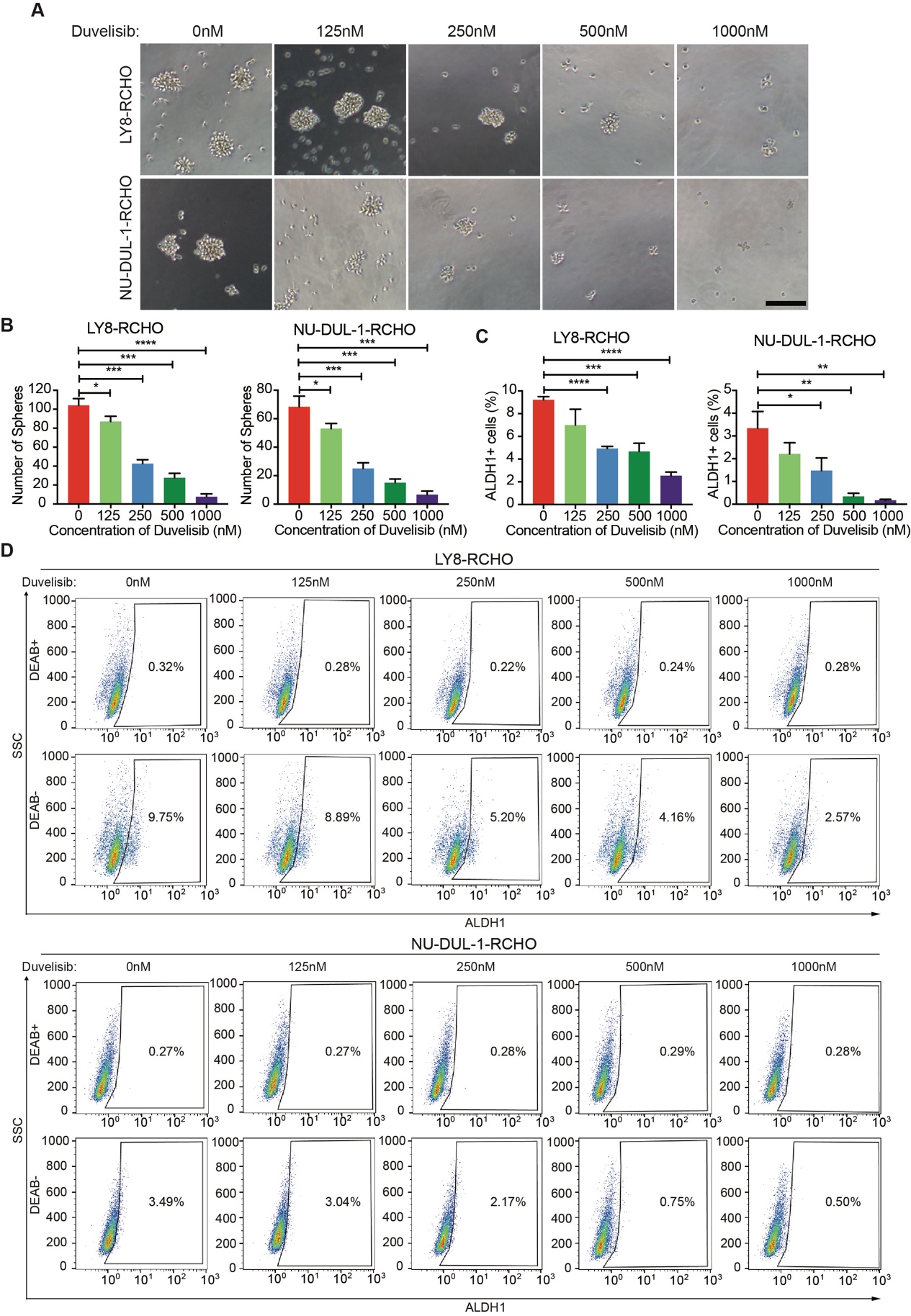

**Fig. S9. Duvelisib treatment reduced sphere-forming capacity of and ALDH1^+^ subpopulation in RCHO-resistant cells.** (**A**-**B**) Duvelisib treatment reduced the sphere-forming capacity of RCHO-resistant DLBCL cells. Representative images (A) and quantitative results (B). Scale bar: 100 µm. (**C**-**D**) Aldefluor assay: duvelisib treatment reduced ALDH1^+^ subpopulation. The RCHO-resistant DLBCL cells were treated by duvelisib for 24 hours before the Aldefluor assay. The DEAB controls were used for gating the ALDH1^+^ cells. Quantitative results (C) and representative images (D). The data are presented as mean ± SD; n=3. **P*<0.05, ***P*<0.01, ****P*<0.001 and *****P*<0.0001.

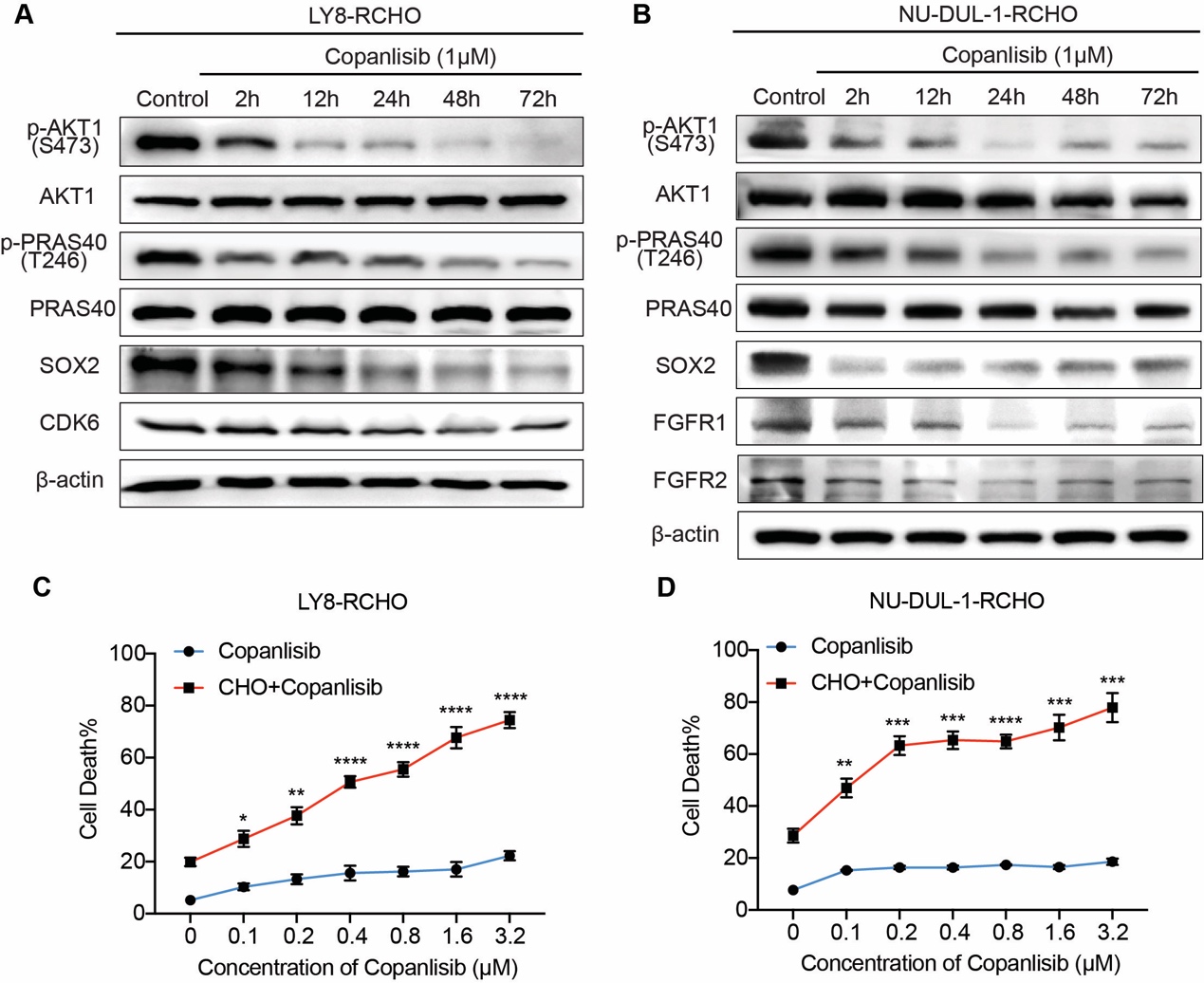

**Fig. S10. Copanlisib reversed the resistance of RCHO-resistant cells to CHO by degrading SOX2.** (**A**-**B**) Copanlisib reduced the expression of SOX2 and CDK6 (A) or FGFR1/2 (B) by suppressing AKT activation in RCHO-resistant LY8 (A) and NU-DUL-1 (B) cells, respectively. (**C**-**D**) CytoTox-Glo cytotoxicity assays: combination of copanlisib with CHO significantly reversed CHO resistance; however, copanlisib alone showed a negligible effect on direct induction of cell death. The cells were treated with copanlisib in the presence or absence of CHO for 48 hours before assays. The data are presented as mean ± SD; n=3. **P*<0.05, ***P*<0.01, ****P*<0.001, and *****P*<0.0001 *vs* CHO alone.

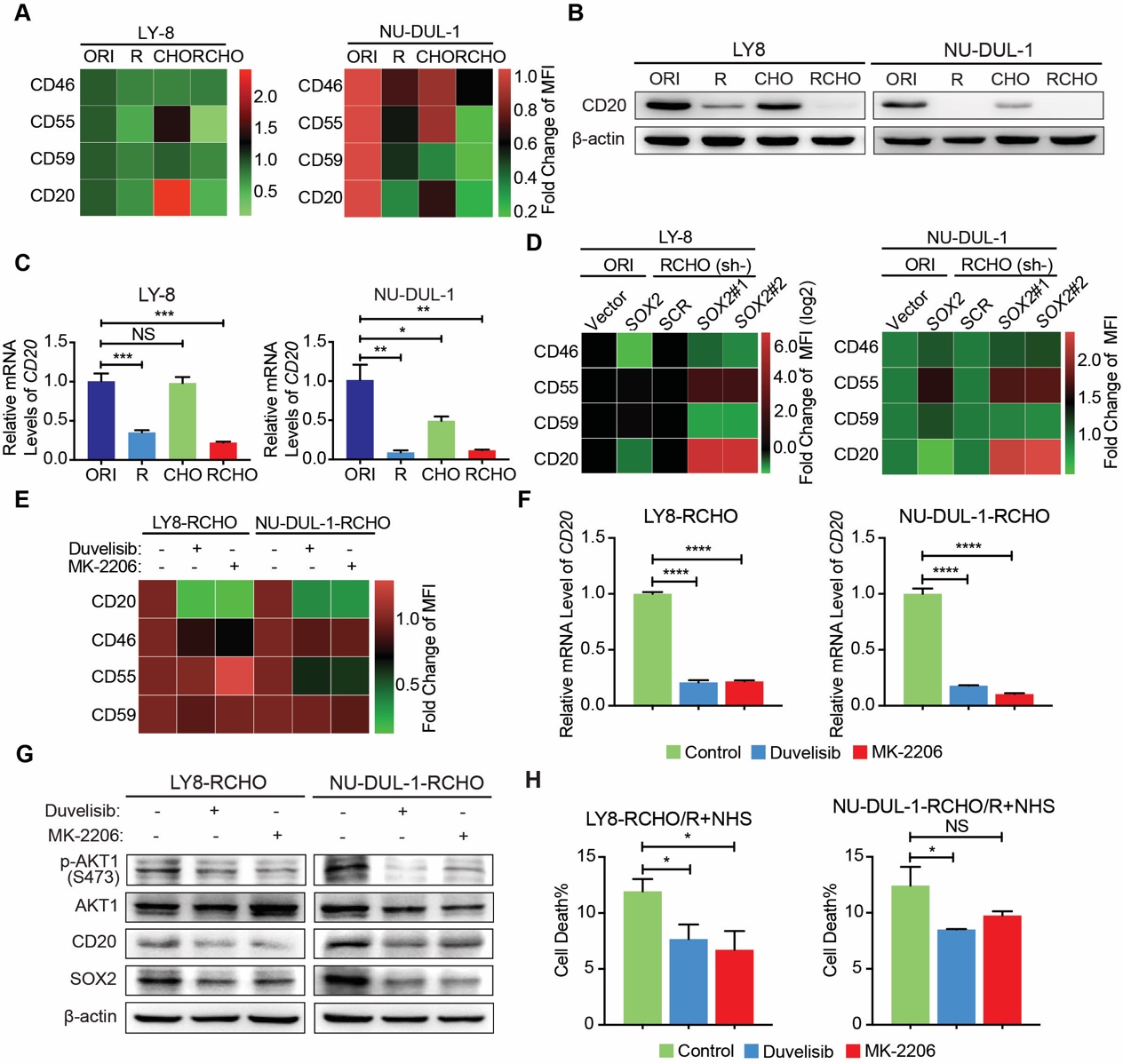

**Fig. S11. The CD20 level was the determining factor for SOX2-induced resistance to R-mediated CDC.** (**A**) FACS analysis for evaluating CD46, CD55, CD59 and CD20 membrane expression in original and resistant DLBCL cells. Quantitative data are shown by heatmaps, with each cell representing mean of three biological repeats. For development of resistance to R-mediated CDC, the membrane levels of mCRPs, namely, CD46, CD55 and CD59, were not elevated, while the membrane CD20 level was significantly reduced in R- and RCHO-resistant DLBCL cells. (**B-C**) CD20 expression clearly declined in resistant cells, especially R- and RCHO-resistant cells, at both the protein (B) and mRNA (C) levels. The data are presented as mean ± SD; n=3. **P*<0.05, ***P*<0.01, and ****P*<0.001. (**D**) FACS analysis for evaluating CD46, CD55, CD59 and CD20 expression in original DLBCL cells with ectopic *SOX2* expression and in *SOX2*-insufficient RCHO-resistant DLBCL cells. Quantitative data are shown by heatmaps (LY8 cells in left panel, and NU-DUL-1 cells in right panel), with each cell representing mean of three biological repeats. For regulation of resistance to R-mediated CDC, ectopic SOX2 expression in original DLBCL cells reduced membrane CD20 levels, while SOX2 insufficiency in RCHO-resistant DLBCL cells increased the membrane CD20 level. However, mCRP expression did not support the role of SOX2 in development of resistance to R-mediated CDC. These results revealed the mechanism by which SOX2 modulated the sensitivity of DLBCL cells to R-mediated CDC by regulating CD20 expression (see Fig. 1K). (**E**) FACS analysis for evaluating CD46, CD55, CD59 and CD20 expression in RCHO-resistant DLBCL cells after duvelisib or MK-2206 treatment. Quantitative data are shown by heatmap, with each cell representing mean of three biological repeats. The expression of CD20, but not mCRPs, markedly decreased after duvelisib or MK-2206 treatment. The discrepancy between PI3K/AKT and SOX2 (Fig. S11D and E) in regulation of CD20 expression suggested that PI3K/AKT regulated CD20 expression via an alternative, SOX2-independent pathway. (**F nad G**) PI3K/AKT inhibition by duvelisib or MK-2206 markedly reduced CD20 expression at both the mRNA (F) and protein (G) levels. (**H**) CDC assays: conversely, PI3K/AKT inhibition by duvelisib or MK-2206 exacerbated resistance to R-mediated CDC in RCHO-resistant DLBCL cells. The data are presented as mean ± SD; n=3. **P*<0.05, and *****P*<0.0001.

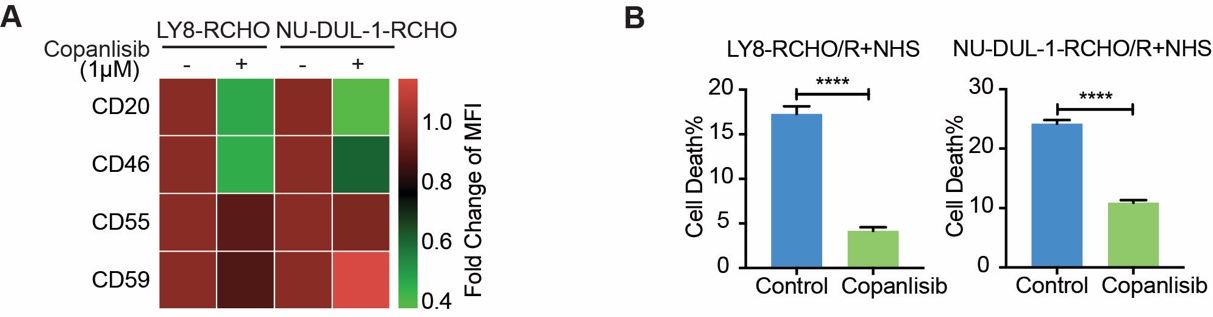

**Fig. S12. Inhibition PI3K/AKT by copanlisib failed to reverse the resistance of RCHO-resistant cells to R-mediated CDC.** (**A**) FACS analysis: CD46, CD55, CD59 and CD20 expression in RCHO-resistant DLBCL cells after copanlisib (1 μM) treatment for 24 hours. Quantitative data are shown by heatmap, with each cell representing mean of three biological repeats. The expression of CD20 markedly decreases after copanlisib treatment. (**B**) CDC assays: PI3K/AKT inhibition by copanlisib instead exacerbated the resistance of RCHO-resistant DLBCL cells to R-mediated CDC. The RCHO-resistant DLBCL cells were pre-treated by copanlisib (1μM) for 24 hours before rituximab (32 μg/mL) and 20% NHS treatment. The data are presented as mean ± SD; n=3. *****P*<0.0001.

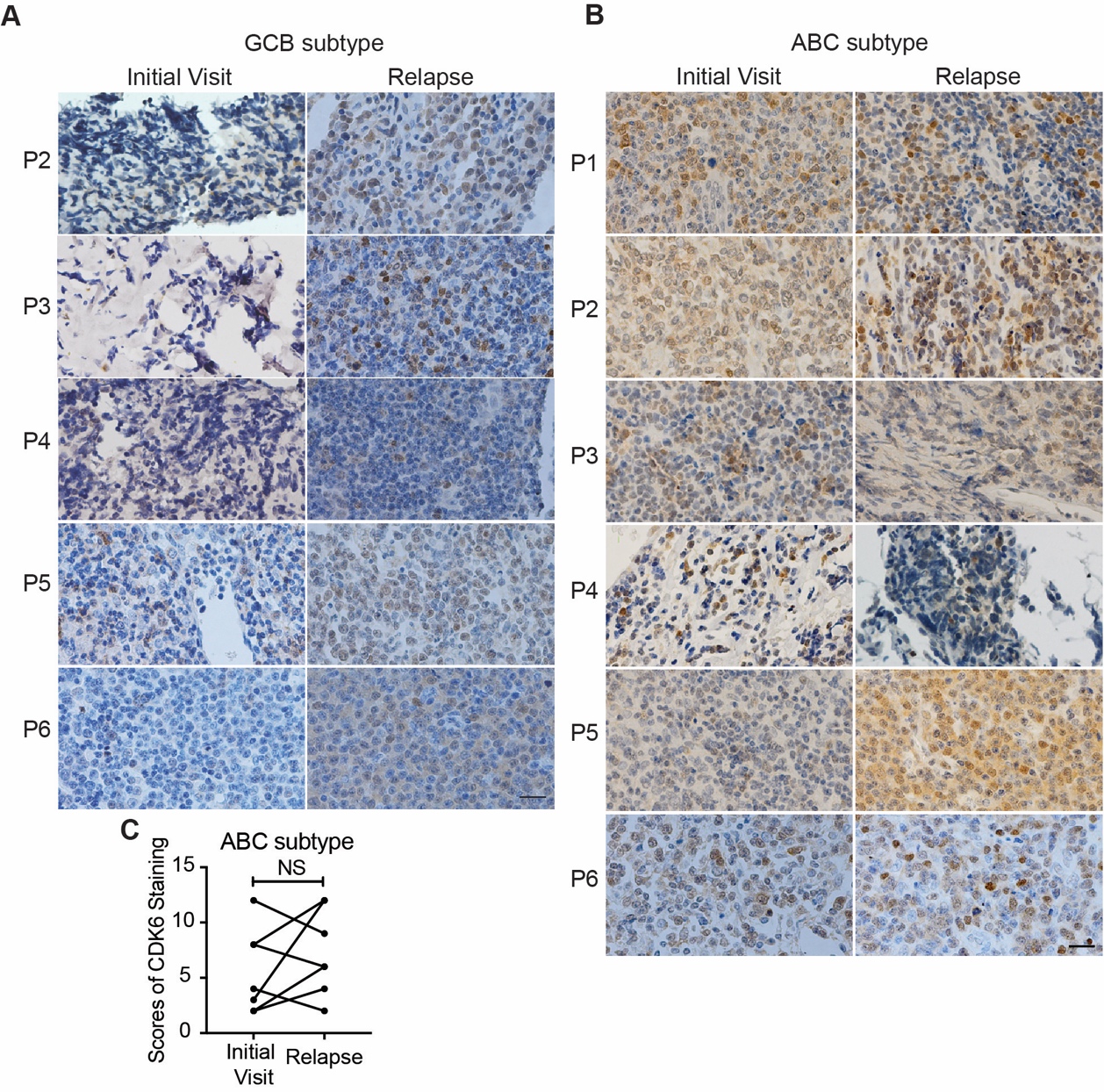

**Fig. S13. CDK6 expression increased only in relapsed GCB but not ABC subtype DLBCL tissues.** (**A**-**C**) CDK6 expression detected by IHC staining significantly increased in relapsed 6 GCB (A) but not 6 ABC (B and C) subtype DLBCL tissues compared with the paired patient tissues from the initial visit. The quantitative score for CDK6 staining of ABC subtype tissues is shown in C and was calculated from B. Image from patient 1 and quantitative results for the GCB subtype are shown in Fig. 4C. P: patient. NS: no significance. Scale bar: 20 μm.

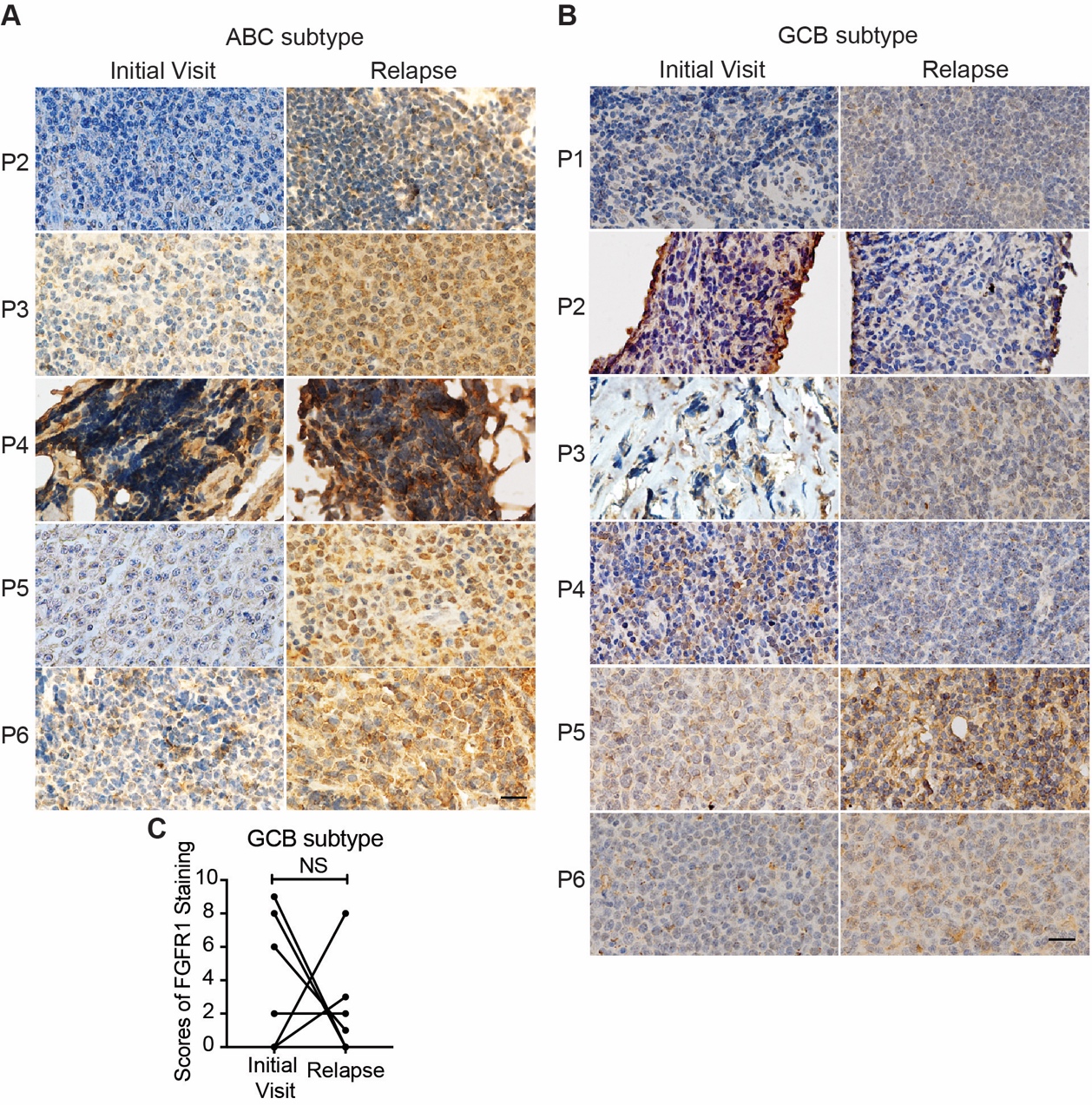

**Fig. S14. FGFR1 expression increased only in relapsed ABC but not GCB subtype DLBCL tissues.** (**A**-**C**) FGFR1 expression determined by IHC staining significantly increased in relapsed 6 ABC (A) but not 6 GCB (B and C) subtype DLBCL tissues compared with the paired patient tissues from the initial visit. The quantitative score for FGFR1 staining of GCB subtype tissues is shown in C and was calculated from B. Image from patient 1 and quantitative results for the ABC subtype are shown in Fig. 4E. P: patient. NS: no significance. Scale bar: 20 μm.

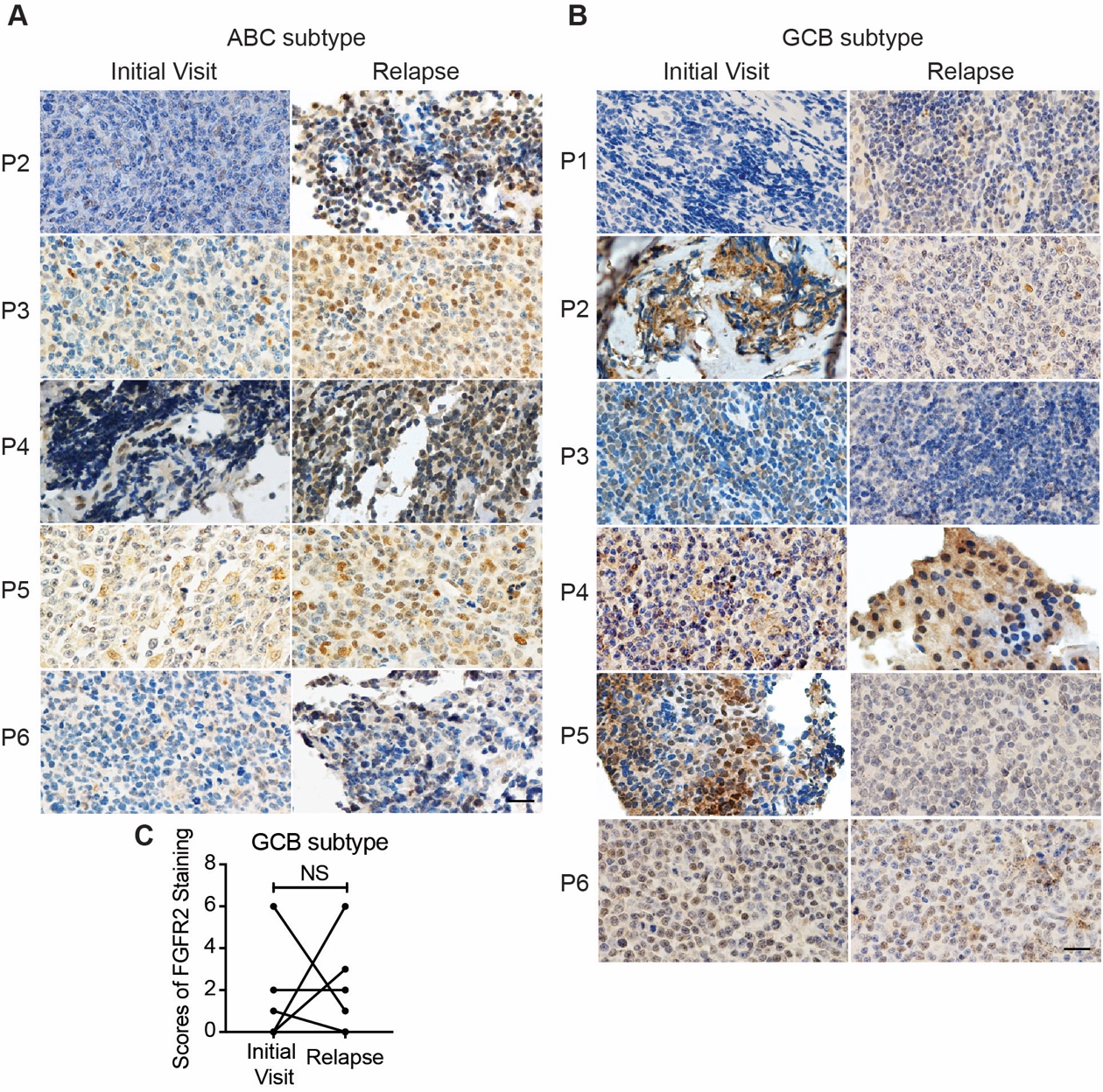

**Fig. S15. FGFR2 expression was elevated only in relapsed ABC but not GCB subtype DLBCL tissues.** (**A**-**C**) FGFR2 expression detected by IHC staining was significantly increased in relapsed 6 ABC (A) but not 6 GCB (B and C) subtype DLBCL tissues compared with the paired patient tissues from the initial visit. The quantitative score for FGFR2 staining of GCB subtype tissues is shown in C and was calculated from B. Image from patient 1 and quantitative results for the ABC subtype are shown in Fig. 4F. P: patient. NS: no significance. Scale bar: 20 μm.

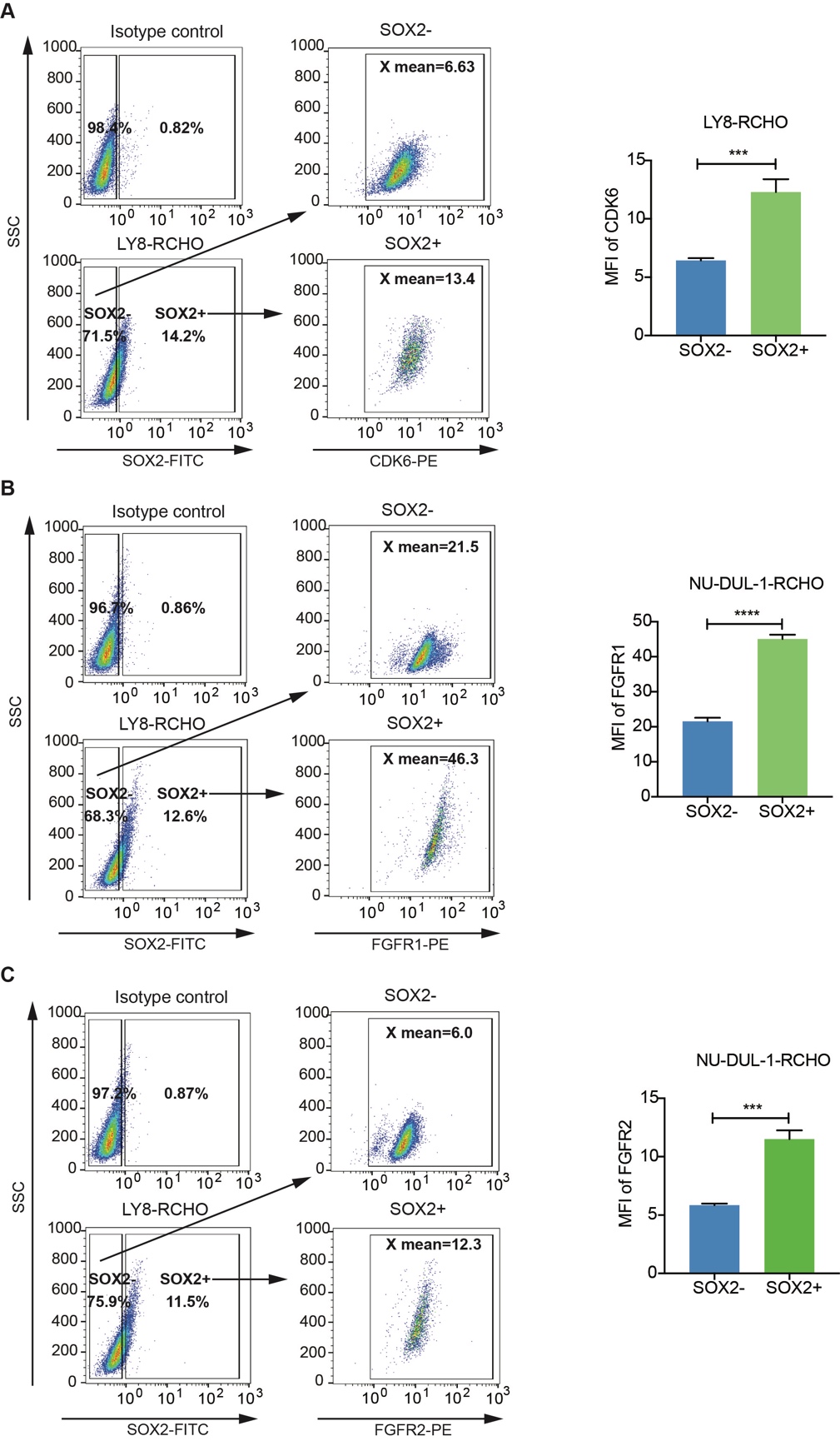

**Fig.S16. SOX2^+^ subpupulation in RCHO-resistant cells showed higher CDK6 or FGFR1/2 level.** (**A**) Representative images (left) for detecting CDK6 levels in SOX2^-^ and SOX2^+^ subpopulations of LY8-RCHO cells. The Fluorochrome-conjugated isotype-specific IgGs were used for gating the SOX2^+^ cells and CDK6^+^ cells. SOX2^+^ subpopulation exhibits higher CDK6 level. The related quantitative results are shown in the right panel. (**B**-**C**) Representative images (left) for detecting FGFR1/2 levels in SOX2^-^ and SOX2^+^ subpopulations of NU-DUL-1-RCHO cells. The Fluorochrome-conjugated isotype-specific IgGs were used for gating the SOX2^+^ cells and FGFR1/2^+^ cells. SOX2^+^ subpopulation exhibits higher FGFR1/2 level. The related quantitative results are shown in the right panel. The data are presented as mean ± SD; n=3. ****P*<0.001, *****P*<0.0001.

**Table S1. The commercial antibodies used in this study.**

| **Antibodies** | **Manufacturers** | **Applications in this study** | **Catalog Number** |
| --- | --- | --- | --- |
| CD34 [ICO-115] | abcam | WB (1:1,000) | ab187282 |
| CD133 | HuaBio Co., Ltd. | WB(1:1,000) | R121101 |
| β-actin (C4) | Santa Cruz Biotechnology | WB (1:1,000) | sc-47778 |
| Sox2 (D6D9) XP Rabbit mAb | Cell Signaling Technology | WB (1:1,000),  IHC (1:200) | 3579 |
| OCT4 [EPR17929] | abcam | WB (1:1,000) | ab181557 |
| Nanog [EPR2027(2)] | abcam | WB (1:1,000) | ab109250 |
| KLF4 [EPR19590] | abcam | WB (1:1,000) | ab215036 |
| c-Myc (D3N8F) Rabbit mAb | Cell Signaling Technology | WB (1:1,000) | 13987 |
| Phospho-Akt (Ser473) (D9E) | Cell Signaling Technology | IHC (1:100)  WB (1:1,000) | 4060 |
| PI3 Kinase p110α (C73F8) | Cell Signaling Technology | WB (1:1,000) | 4249 |
| PI3 Kinase p110β (C33D4) | Cell Signaling Technology | WB (1:1,000) | 3011 |
| PI3 Kinase p110γ (D55D5) | Cell Signaling Technology | WB (1:1,000) | 5405 |
| PI3 Kinase p110 δ (D1Q7R) | Cell Signaling Technology | WB (1:1,000) | 34050 |
| PI3 Kinase p85 (19H8) | Cell Signaling Technology | WB (1:1,000) | 4257 |
| Akt1 (C73H10) | Cell Signaling Technology | WB (1:1,000) | 2938 |
| Ubiquitin (P4D1) | Cell Signaling Technology | WB (1:1,000) | 3936 |
| Phospho-FAK (Tyr397) (D20B1) | Cell Signaling Technology | WB (1:1,000) | 8556 |
| FAK (D2R2E) | Cell Signaling Technology | WB (1:1,000) | 13009 |
| Integrin α1(A-9) | Santa Cruz Biotechnology | WB (1:500) | sc-271034 |
| Integrin beta 5 | abcam | WB (1:1,000) | ab15459 |
| Phospho-Akt1 (Ser473) (D7F10) | Cell Signaling Technology | WB (1:1,000) | 9018 |
| CD79a [EPR3619] | abcam | WB (1:1,000) | ab133483 |
| p-Lyn (Y 396) | abcam | WB (1:1,000) | ab226778 |
| Lyn (C13F9) | Cell Signaling Technology | WB (1:1,000) | 2796 |
| SHP-1 [EPR5519] | abcam | WB (1:1,000) | ab124942 |
| Phospho-SHP-1 (Tyr564) (D11G5) | Cell Signaling Technology | WB (1:1,000) | 8849 |
| TCF3/TCF7L1 (D15G11) | Cell Signaling Technology | WB (1:1,000) | 2883 |
| Phospho-Syk (Tyr525/526) (C87C1) | Cell Signaling Technology | WB (1:1,000) | 2710 |
| Syk (N-19) | Santa Cruz Biotechnology | WB (1:500) | sc-1077 |
| CCR7 [E75] | abcam | WB (1:500) | ab32075 |
| p-Src(Y418) [EP503Y] | abcam | WB (1:1,000) | ab40660 |
| Src [EPR5496] | abcam | WB (1:1,000) | ab109381 |
| CDK6 [EPR4515] | abcam | WB (1:1,000)  IHC (1:100) | ab124821 |
| FGFR1 | abcam | WB (1:1,000)  IHC (1:400) | ab10646 |
| FGFR2 (C17) | Santa Cruz Biotechnology | WB (1:1,000)  IHC (1:200) | sc-122 |
| CD20 (L26) | Santa Cruz Biotechnology | WB (1:1,000) | sc-58985 |
| PE anti-human CD55 (IA10) | BD Pharmingen | FACS (20 µL/test) | 555694 |
| PE anti-Human CD59 (H19) | BioLegend | FACS (5 µL/test) | 304708 |
| APC anti-human CD46 (TRA-2-10) | BioLegend | FACS (5 µL/test) | 352405 |
| APC anti-human CD20 (2H7) | BioLegend | FACS (5 µL/test) | 302310 |
| APC anti-human CD133 | BioLegend | FACS (5 µL/test) | 372806 |
| Human TruStain FcX^TM^ (Fc Receptor Blocking Solution) | BioLegend | FACS (5 µL/test) | 422301 |
| FITC Goat anti-mouse IgG (minimal x-reactivity) | BioLegend | FACS (1 µg/test) | 405305 |
| PE Donkey anti-rabbit IgG (minimal x-reactivity) | BioLegend | FACS (0.125 µg/test) | 406421 |
| APC Goat anti-mouse IgG (minimal x-reactivity) | BioLegend | FACS (0.125 µg/test) | 405308 |
| Phospho-PRAS40 (Thr246) (C77D7) Rabbit mAb | Cell Signaling Technology | WB (1:1,000) | 2997 |
| PRAS40 (D23C7) XP® Rabbit mAb | Cell Signaling  Technology | WB (1:1,000) | 2691 |
| PAX5 (D7H5X) XP® Rabbit mAb | Cell Signaling  Technology | WB (1:1,000) | 12709 |
| Sox2 (D9B8N) Rabbit mAb | Cell Signaling  Technology | IP (1:100)  FACS (1:100) | 23064 |
| Sox2 (L1D6A2) Mouse mAb | Cell Signaling  Technology | FACS (1:200) | 4900 |
| Goat anti-mouse IgG-HRP | Santa Cruz Biotechnology | WB (1:10,000) | sc-2005th |
| Goat anti-rabbit IgG-HRP | Santa Cruz Biotechnology | WB (1:10,000) | sc-2004 |
| Peroxidase-conjugated affinipure goat anti-rabbit IgG H&L | Proteintech Group Inc. | IHC (1:200) | SA00001-2 |
| Normal mouse IgG | Santa Cruz Biotechnology | 2 µg/test | sc-2025 |

**Table S2. The sequences of primers and shRNA used in this study.**

| **Primers** | **5' to 3'** |
| --- | --- |
| *SOX2* qRT-PCR forward primer | GCCGAGTGGAAACTTTTGTCG |
| *SOX2* qRT-PCR reverse primer | GGCAGCGTGTACTTATCCTTCT |
| *ITGA1* qRT-PCR forward primer | CGCTGCTGCGTATCATTCAA |
| *ITGA1* qRT-PCR reverse primer | GGCCAACTAACGGAGAACCA |
| *ITGA2* qRT-PCR forward primer | AGTGGCTTTCCTGAGAACCG |
| *ITGA2* qRT-PCR reverse primer | GATCAAGCCGAGGCTCATGT |
| *ITGA2B* qRT-PCR forward primer | AACACCCTGAGCCGCATTTA |
| *ITGA2B* qRT-PCR reverse primer | AATAATAGCCGCCAGGAGCC |
| *ITGA3* qRT-PCR forward primer | CCCTATTCCTCCGAACCAGC |
| *ITGA3* qRT-PCR reverse primer | CGCCTTCTGCCTCTTAGCTT |
| *ITGA4* qRT-PCR forward primer | CAACGAAAGAACTGGACGGC |
| *ITGA4* qRT-PCR reverse primer | TAGAAGCGGAGCTGTGTGAC |
| *ITGA5* qRT-PCR forward primer | ACATCTGTGTGCCTGACCTG |
| *ITGA5* qRT-PCR reverse primer | CTCACCCACATTCTGGGCAT |
| *ITGA6* qRT-PCR forward primer | TTCGGGAGTACCTTGGTGGA |
| *ITGA6* qRT-PCR reverse primer | AGAGCGTTTAAAGAATCCACACT |
| *ITGA7* qRT-PCR forward primer | ATTTCCCTTGCATTCGCTGG |
| *ITGA7* qRT-PCR reverse primer | CGTCCAGATTGAAGGCGACA |
| *ITGA8* qRT-PCR forward primer | CTTATTGTGGGAGGACCTGGG |
| *ITGA8* qRT-PCR reverse primer | GTAAACTCCCCAGCAGCAACT |
| *ITGA9* qRT-PCR forward primer | GGCTGTGTTTAAGTGCCGTG |
| *ITGA9* qRT-PCR reverse primer | ATCCACTCATCATCGCGGTC |
| *ITGA10* qRT-PCR forward primer | GATCAGGCATGGAACTCCCC |
| *ITGA10* qRT-PCR reverse primer | ACAGGGCAGCGATAAACGTC |
| *ITGA11* qRT-PCR forward primer | TGGCCAGGGTTCACGGAC |
| *ITGA11* qRT-PCR reverse primer | CCGTGGATCACTGGACACTT |
| *ITGAD* qRT-PCR forward primer | CATGAGATTCAGCCCTGTGGA |
| *ITGAD* qRT-PCR reverse primer | TCCAGTGCCAGATCAAACCT |
| *ITGAE* qRT-PCR forward primer | CTCAAGTGGGGAGTGTCACC |
| *ITGAE* qRT-PCR reverse primer | CAACACCCTGCATTTTGGGG |
| *ITGAL* qRT-PCR forward primer | CTGTAAGAGGCCAAAGGGCA |
| *ITGAL* qRT-PCR reverse primer | GCGCGAAGAAAAAGAACCCA |
| *ITGAM* qRT-PCR forward primer | GCCAGGACCTCACAATGGAT |
| *ITGAM* qRT-PCR reverse primer | CTTGCCACTTCCCTGGGATT |
| *ITGAV* qRT-PCR forward primer | TCCCATCAGTGGTTTGGAGC |
| *ITGAV* qRT-PCR reverse primer | TGATCTACATGGAGCATACTCAACA |
| *ITGAX* qRT-PCR forward primer | TGGCTTCTTCAAGCGTCAGT |
| *ITGAX* qRT-PCR reverse primer | GAGGGTAATGGGGAGTGGGC |
| *ITGB1* qRT-PCR forward primer | GCCGCGCGGAAAAGATGAAT |
| *ITGB1* qRT-PCR reverse primer | GAATTTGTGCACCACCCACAA |
| *ITGB2* qRT-PCR forward primer | GTGTCAGGACTTTACGACCCG |
| *ITGB2* qRT-PCR reverse primer | ACTCCTGAGAGAGGACGCA |
| *ITGB3* qRT-PCR forward primer | ACCAGTAACCTGCGGATTGG |
| *ITGB3* qRT-PCR reverse primer | CTCATTGAAGCGGGTCACCT |
| *ITGB4* qRT-PCR forward primer | AAGGGCAACATCCATCTGAAAC |
| *ITGB4* qRT-PCR reverse primer | GCTGACCGCACCTCTTTTTG |
| *ITGB5* qRT-PCR forward primer | ATACCTGGAACAACGGTGGAG |
| *ITGB5* qRT-PCR reverse primer | GGCTGATCCCAGACTGACAA |
| *ITGB6* qRT-PCR forward primer | ATCGGTCTGCACAGCAAGAA |
| *ITGB6* qRT-PCR reverse primer | CAGGCACACTGAGGTCCAAT |
| *ITGB7* qRT-PCR forward primer | GGCTCTTCTACCACTACGGC |
| *ITGB7* qRT-PCR reverse primer | CATTCTGTGGCATCCCCTGT |
| *ITGB8* qRT-PCR forward primer | AGCTGCAACTAATGGTGTTGG |
| *ITGB8* qRT-PCR reverse primer | AACCCAAACAAAGCCCGAAG |
| *CCR7* qRT-PCR forward primer | ATGCCTGTGTCAAGATGAGGTCA |
| *CCR7* qRT-PCR reverse primer | AAGTTCCGCACGTCCTTCTT |
| *CD79A* qRT-PCR forward primer | AGAACGAGAAGCTCGGGTTG |
| *CD79A* qRT-PCR reverse primer | CCCCGGGAGATGTCCTCATA |
| *CD20* qRT-PCR forward primer | AACTCAGCAGTAGGCCTTGC |
| *CD20* qRT-PCR reverse primer | GCAGTCTTACCTTGTGTCATGC |
| *Firefly Luciferase* CDS  forward primer | ATGGAAGACGCCAAAAACATAAAG |
| *Firefly Luciferase* CDS  reverse primer | TTACACGGCGATCTTTCCGCCCTT |
| *myr-AKT1* CDS  forward primer | GCGAATTCATGGGCTGTGGCTGCAGCTCACACCCGGAAGATGACATGAGCGACGTGGCTATTGTG |
| *myr-AKT1* CDS  reverse primer | GCGGATCCTCACTTGTCATCGTCGTCCTTGTAATCGGCCGTGCCGCTGGCCGAG |
| human *OCT4* CDS  forward primer | ATGGCGGGACACCTGGCT |
| human *OCT4* CDS  reverse primer | TCAGTTTGAATGCATGGGAGA |
| *SOX2* shRNA#1 forward primer | AATTAGGAGCACCCGGATTATAAATCTCGAGATTT  ATAATCCGGGTGCTCCTTTTTTTTAT |
| *SOX2* shRNA#1 reverse primer | AAAAAAAAGGAGCACCCGGATTATAAATCTCGAGA  TTTATAATCC GGGTGCTCCT |
| *SOX2* shRNA#2 forward primer | AATTGGTTGACACCGTTGGTAATTTCTCGAGAAAT  ACCAACGGTGTCAACCTTTTTTTAT |
| *SOX2* shRNA#2 reverse primer | AAAAAAAGGTTGACACCGTTGGTAATTTCTCGAGA  AATTACCAACGGTGTCAACC |
| *ITGA1* shRNA#1 forward primer | AATTTCTGGAGATGTGCTCTATATTCTCGAGAATATAGAGCACATCTCCAGATTTTTTTAT |
| *ITGA1* shRNA#1 reverse primer | AAAAAAATCTGGAGATGTGCTCTATATTCTCGAGAATATAGAGCACATCTCCAGA |
| *ITGA1* shRNA#2 forward primer | AATTACTCTGGGAGAAAGAATATTTCTCGAGAAATATTCTTTCTCCCAGAGTTTTTTTTAT |
| *ITGA1* shRNA#2 reverse primer | AAAAAAAACTCTGGGAGAAAGAATATTTCTCGAGAAATATTCTTTCTCCCAGAGT |
| *ITGB5* shRNA#1 forward primer | AATTGGATTGGAAGTAAAGATTAAACTCGAGTTTAATCTTTACTTCCAATCCTTTTTTTAT |
| *ITGB5* shRNA#1 reverse primer | AAAAAAAGGATTGGAAGTAAAGATTAAACTCGAGTTTAATCTTTACTTCCAATCC |
| *ITGB5* shRNA#2 forward primer | AATTTTCCTTCATCTTGTGTAAATACTCGAGTATTTACACAAGATGAAGGAATTTTTTTAT |
| *ITGB5* shRNA#2 reverse primer | AAAAAAATTCCTTCATCTTGTGTAAATACTCGAGTATTTACACAAGATGAAGGAA |
| *CD79A* shRNA#1 forward primer | AATTACTTCCAATGCCCGCACAATACTCGAGTATTGTGCGGGCATTGGAAGTTTTTTTTAT |
| *CD79A* shRNA#1 reverse primer | AAAAAAAACTTCCAATGCCCGCACAATACTCGAGTATTGTGCGGGCATTGGAAGT |
| *CD79A* shRNA#2 forward primer | AATTCAGGGCCACTTAGTGATAATACTCGAGTATTATCACTAAGTGGCCCTGTTTTTTTAT |
| *CD79A* shRNA#2 reverse primer | AAAAAAACAGGGCCACTTAGTGATAATACTCGAGTATTATCACTAAGTGGCCCTG |
| *CCR7* shRNA#1 forward primer | AATTCGAGCTTGTTCTTTGTTCTTTCTCGAGAAAGAACAAAGAACAAGCTCGTTTTTTTAT |
| *CCR7* shRNA#1 reverse primer | AAAAAAACGAGCTTGTTCTTTGTTCTTTCTCGAGAAAGAACAAAGAACAAGCTCG |
| *CCR7* shRNA#2 forward primer | AATTCCCTTTCTTGTACGCCTTCATCTCGAGATGAAGGCGTACAAGAAAGGGTTTTTTTAT |
| *CCR7* shRNA#2 reverse primer | AAAAAAACCCTTTCTTGTACGCCTTCATCTCGAGATGAAGGCGTACAAGAAAGGG |
| scramble shRNA forward primer | AATTCCTAAGGTTAAGTCGCCCTCGCTCGAGCGAGGGCGACTTAACCTTAGGTTTTTTT |
| scramble shRNA reverse primer | AAAAAAACCTAAGGTTAAGTCGCCCTCGCTCGAGCGAGGGCGACTTAACCTTAGG |

**Table S3. Methods of drug delivery in xenograft model.**

| **Drug** | **Dosage (mg/kg)** | **Administration Day**  **(Days after tumor implantation)** | **Administration Route** |
| --- | --- | --- | --- |
| Rituximab (R) | 118.4 | 8 | intraperitoneal injection |
| Cyclophosphamide (C) | 235.6 | 8 | intraperitoneal injection |
| Doxorubicin (H) | 16 | 8 | intraperitoneal injection |
| Vincristine (O) | 0.444 | 8 | intraperitoneal injection |
| Prednisolone (P) | 12.3 | 8, 9, 10, 11, 12 | intraperitoneal injection |
| Duvelisib | 5.18 | 8, 10, 12, 13, 15, 17, 19, 21, 23, 25, 27 | gavage |
| Abemaciclib | 41.1 | 8, 10, 12, 13, 15, 17, 19, 21, 23, 25, 27 | gavage |
| AZD4547 | 32.8 | 8, 9, 10, 11, 12, 13, 14, 15, 16, 17, 18, 19, 20, 21 | gavage |
